# Root Cap-Specific Transcriptomics Identify the NAC Transcription Factor SOMBRERO as a Versatile Regulator of Auxin Gradients

**DOI:** 10.1101/2025.01.22.634243

**Authors:** Pengfei Wang, Zhongyuan Liu, Yuling Zhou, Junxian He, Byung-Ho Kang

## Abstract

Auxin gradients in the root cap play a crucial role in the root cap development and root tropism. In the central root cap, meristematic cells in the columella initial differentiate into gravity-sensing columella cell, and mature into border-like cells that eventually separate from the root cap. Golgi stacks exhibit distinct ultrastructural features in each cell type across the central root cap, serving as effective cell markers. This cell type-specific Golgi remodeling was inhibited in *smb-3* mutant root cap. When we carried out transcriptomic analyses of isolated root cap from Col-0 and *smb-3*, transcript levels of numerous auxin-related genes, including those involved in auxin synthesis, transport, and signaling, were significantly altered in *smb-3* mutant root caps. The auxin gradient was suppressed within the central root cap of *smb-3* mutant lines and they displayed decreased sensitivity to N-1-naphthylphthalamic acid (NPA) in comparison to Col-0. Furthermore, we demonstrated that the SMB binds to the promoter regions of auxin-related genes displaying large changes in *smb-3* root cap cells. Our findings suggest that SMB functions as a versatile transcription factor that regulates the expression of genes critical for local auxin distribution and responses in the root cap, thereby coordinating root cap cell differentiation and controlling root growth directions in response to external stimuli.

## INTRODUCTION

Auxin is an essential phytohormone that regulates various aspects of plant growth and development, including cell elongation, division, and differentiation^1–4^. One of the key features of auxin is its ability to form concentration gradients within plant tissues to control tissue patterning and organ formation^5–11^. The root cap illustrates the significance of local auxin gradient in the plants’ responses to environmental and developmental cues. The dynamic distribution of auxin is critical for plant roots to respond to environmental stimuli^12–17^. It was shown that asymmetric auxin distribution is also involved in the continuous cell renewal in the root cap, maintaining its size^10^.

Auxin gradient is set up via its polar transport by specialized carrier proteins such as PIN-FORMED (PIN) proteins, which facilitate the directional flow of auxin^18–20^. PIN3, PIN4 and PIN7 are auxin efflux carriers, expressed in the central root cap and facilitate auxin distribution within the root cap^8,10,21^. The transporter are required for gravity-sensing by the root cap and their distribution within the cells is adjusted according to the gravity vector direction for reorienting auxin flow^22^. N-1-naphthylphthalamic acid (NPA) is an herbicides that binds to PIN proteins, inhibiting auxin transport. NPA has been employed in plant auxin research to monitor consequences of inhibited auxin polar transport^23,24^. NPA disrupts the auxin gradient in the root cap and induces agravitropic growth^25,26^. In addition to directional auxin flow, auxin biosynthesis in the root by TAA1 and its close homologue, TAR2, also contribute to root cap development^27,28^.

The auxin gradient leads to uneven activation of Auxin Response factor 10 (ARF10) and 16 (ARF16) within the root cap to control cell division and differentiation programs. Expression of the two ARFs is regulated by microRNA160 (miR160) and a non-canonical AUX/IAA protein, IAA33, interacts with ARF10 and ARF16, suggesting a tight regulation of auxin signaling in the root cap^29–31^. However, our understanding is limited concerning how these multiple regulatory networks involving auxin synthesis, polar transport, and signal transduction pathways are orchestrated in the root cap where new cells are produced in the center and terminally differentiated cells are lost from the surface.

Sombrero (SMB) is a member of the NAC (NAM, ATAF1/2, and CUC2) family of transcription factors, which play diverse roles in plant development and stress responses^32–37^. SMB is specifically expressed in the root cap, where it forms a feedback loop with another transcription factor, FEZ, to regulate periclinal cell division in the lateral root cap^33^. SMB also activates programmed cell death in the lateral root cap^38^. Inactivation of SMB disrupts this regulatory loop and inhibits cell death, resulting in disorganization of the root cap architecture and additional cells in the lateral root cap^38,39^. Furthermore, SMB mediates auxin transport and response within the lateral root cap, promoting the plant’s avoidance from high salt concentrations, a process known as halotropism^32^. SMB expression in the central root cap is as high as in the lateral root cap. SMB’s functions within the central root cap, especially as to auxin signaling, have not been characterized.

The cell differentiation program within the central root cap of Arabidopsis is clearly illustrated in electron micrographs of high-pressure frozen root samples^40–42^. This program begins with the dividing cells of the columella initial, which sequentially differentiate into gravity-sensing columella cells. The cells further develop into peripheral cells, where mucilage synthesis is activated, and ultimately into border-like cells (BLCs) that are shed from the root cap^43,44^. The root cap is under cell flux as new cells arise in the columella initial and BLCs are lost from the root cap surface. Each of these four cell types exhibits distinct changes in their organelle shapes, but Golgi stacks display the most pronounced ultrastructural characteristics, indicating cell differentiation status.

In the current study, we have demonstrated that SMB acts as a versatile regulator that involved in auxin synthesis, transport, and response in the root cap to maintain the auxin response gradient. The auxin response gradient and cell differentiation were severely affected in the *smb-3* central root cap. Further analysis revealed that SMB binds directly to the promoter regions of these auxin-related genes to modulate their transcription. Inactivation of SMB reduces the capacity for auxin redistribution from the central root cap and compromises the ability to generate asymmetric auxin distribution in the lateral root cap, leading to significantly reduced sensitivity to external stimulus NPA.

## RESULTS

### The root cap-specific NAC TF SMB is required for root cap maturation and Golgi remodeling

Golgi stacks in the central columella cells undergo morphological transformations that parallel the differentiation of root cap cells from the columella initials to the outermost BLCs (Figures 1A-1C and S1D). Since the root cap organization is disrupted in *smb-3* mutants (Figures 1E, 1H, S1A and S1B), we first examined Golgi stacks in Col-0 root cap cells using quantitative electron microscopy and immunogold labeling. The sizes (volume and surface area) of Golgi stacks increase throughout the differentiation process, peaking when columella cells mature into BLCs (Figures 1B, 1C and S1C). In these BLC Golgi stacks, *trans*-Golgi cisternae had swollen margins, which were labeled by the LM8 antibody detecting xylogalacturonan (Figures 1B, 1C, 1F, 1G, and S2). These findings agree with our previous observation in alfalfa and maize border cells^45,46^. Interestingly, as BLCs became highly vacuolated, the Golgi stacks lost their hypertrophied margins, exhibited an increased number of cisternae, and became deformed as the vacuoles further expanded during the programmed cell death (PCD) of BLCs (Figures 1B-1D and S1G). Given the cell-type specific Golgi architectures in the root cap, we examined the Golgi stacks in *smb-3* root cap cells. Golgi stacks in the *smb-3* root cap resembled those of columella cells in the Col-0 root cap. *smb-3* BLCs had Golgi stacks devoid of peripheral swellings in their *trans* cisternae (Figures 1H-1J, S1E and S1F). Immunogold labeling with LM8 revealed that xylogalacturonan was rarely produced by Golgi stacks in *smb-3* BLCs (Figures 1J and 1K). Given the link between the Golgi remodeling and cell differentiation in the root cap, these results suggest that SMB is crucial for the sequential root cap cell maturation program.

**Figure 1.**
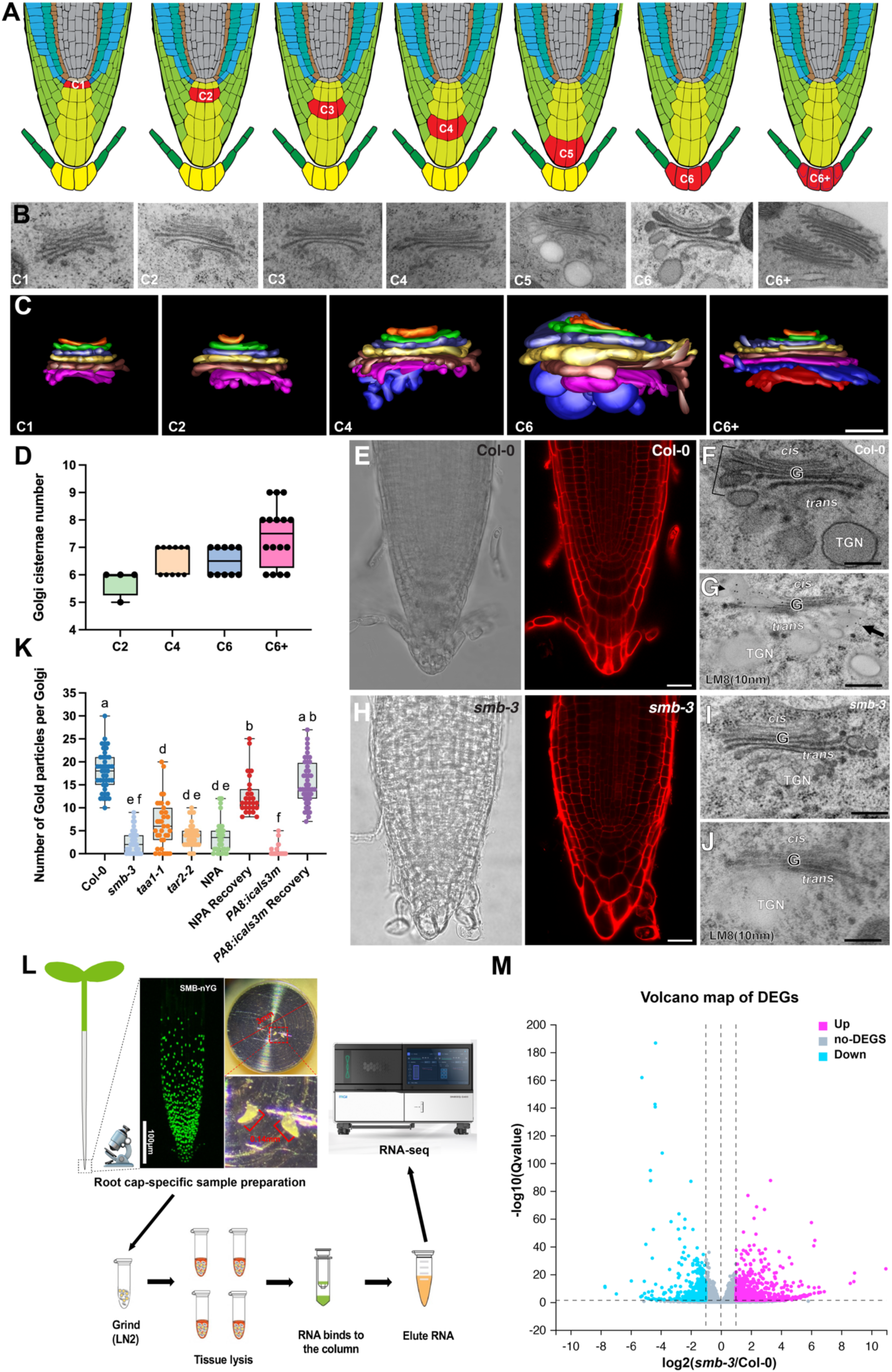
SMB is required for root cap maturation and Golgi remodeling in *Arabidopsis* root cap. (A) Organization of the Arabidopsis root tip. The color-coded cell types in a longitudinal section through the root cap are shown. The cells highlighted in red mark the transition from columella stem cells to border-like cells in the central root cap. The labeled cell types are: C1(columella stem cells), C2(early columella cells), C4(late columella cells), C6(border-like cells (BLCs)), and C6+ (highly vacuolated BLCs). (B) Electron micrographs of Golgi stacks from columella stem cells to highly vacuolated BLCs, as shown in A. (C) Tomographic model of the Golgi stack depicted in B. (D) Statistical analysis of Golgi cisternae number from C2 to C6+. (E-J) Light microscopy images of propidium iodide (PI) stained root tip of Col-0(E) and *smb-3*(H); Electron micrographs showing the ultrastructure of Golgi stacks in BLCs of Col-0(F) and *smb-3*(I); Electron micrographs showing Golgi labeled with an anti-XGA antibody LM8 in BLCs of Col-0(G) and *smb-3*(J). (K) Statistical analysis of XGA-specific gold particles across different mutants or treatments. (L) Experimental design and workflow for root cap-specific sample isolation and RNA enrichment. (M) Volcano plot displaying differentially expressed genes (DEGs) between Col-0 and *smb-3*, based on root cap-specific RNA-seq analysis. The X-axis represents log2 transformed fold changes, while the Y axis represents the significance value after conversion to -log10. Magenta indicates upregulated DEGs, blue indicates downregulated DEGs, and gray indicates non-DEGs. Data are presented as mean ± SD. Alphabets indicate significant differences(p<0.05, one-way ANOVA by Tukey’s test). Scale bars: C, F-G, I-J: 250 nm; E and H: 50 μm. See also Figures S1 and S2

### A root cap-specific RNA-seq reveals that SMB is involved in auxin regulation within the root cap

We carried out comparative RNA-seq analyses of Col-0 and *smb-3* to identify candidate genes that regulated by SMB. For minimizing noises from non-root cap cells, we dissected out samples from 0.15-0.25mm length root tips and prepared RNA samples from the samples (Figure 1L). As a control, RNA samples from root tissues were also isolated. Our root cap-specific transcriptome analysis revealed approximately 3000 differentially expressed genes (DEGs) between Col-0 and *smb-3* (Figure 1M, log_2_FC>0.5, FC>1.414). Gene Ontology (GO) term analyses of these DEGs highlighted a cluster of auxin-related genes (Figures 2A and 2B), suggesting that SMB may control genes involved in auxin-related processes in the root cap. Among auxin-related genes, we firstly focused on the auxin signalling transduction pathway, encompassing genes responsible for auxin response factors (e.g., *ARF10* and *ARF16*) and their regulators (e.g., *IAA33* and *miR160*). The auxin-related gene clusters included auxin homeostasis such as auxin biosynthesis through tryptophan-dependent pathway (TAA1/TARs) and auxin transport via PIN efflux transporters in the root cap (Figure 2C). Figure S3F shows representative genes that up- or down-regulated in the *smb-3* mutant, which could correspond to direct targets of SMB. qRT-PCR analyses of the genes verified their RNA-seq results and supported that SMB may govern multiple auxin-related pathways in the root cap (Figure S3G). Moreover, our root cap-specific RNA-seq allowed us to identify novel, potentially root cap-specific expressed genes (Figures S3C and S3D; Table S1) and widely expressed genes that are specifically regulated by root cap transcription factor SMB (Figures S3A and S3B). Interestingly, some genes that are generally not or scarcely expressed in the root cap but are highly expressed in the root cap when SMB is inactive, providing evidence that SMB could be a root cap identity(Figure S3H; Table S2).

**Figure 2.**
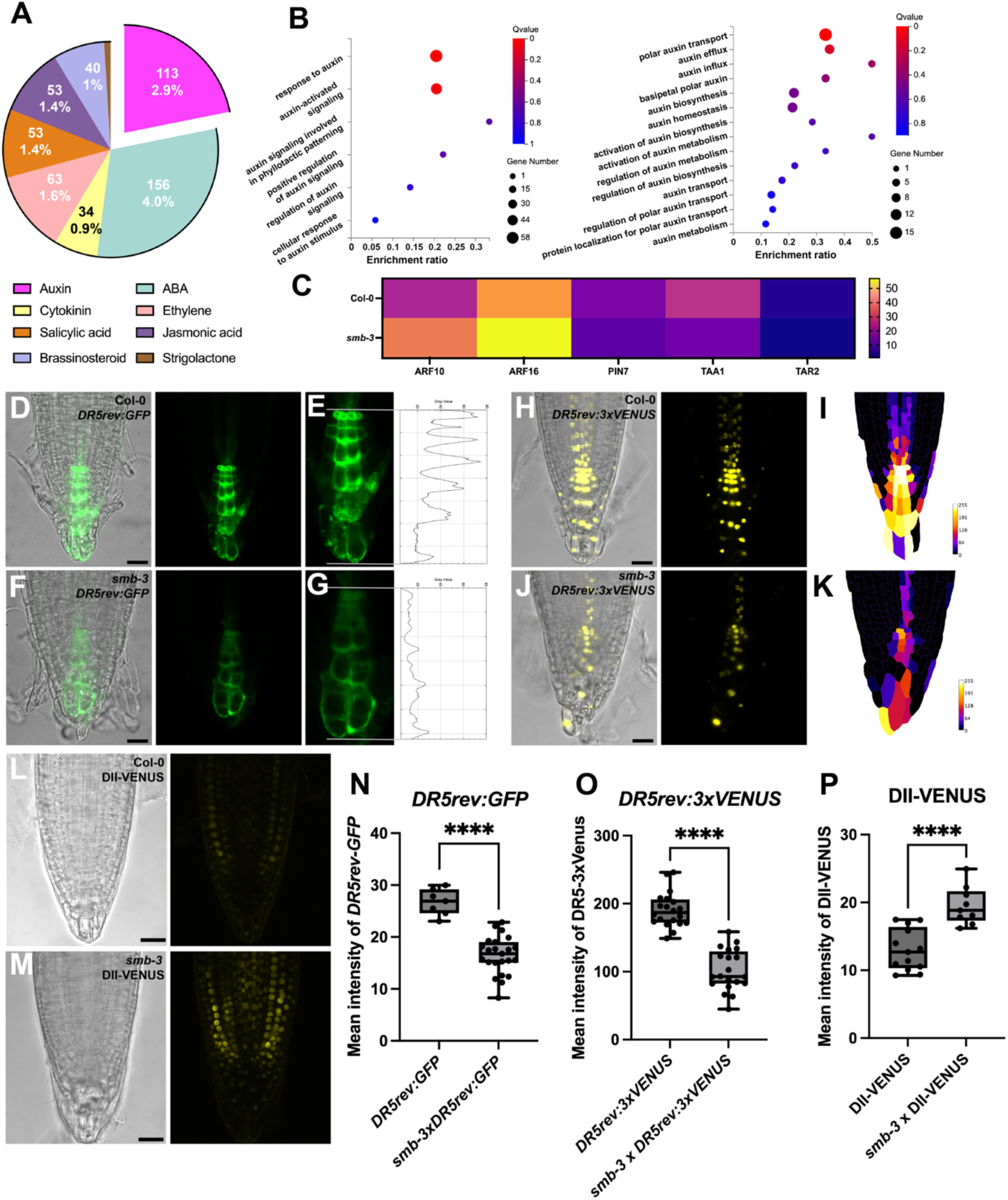
A local auxin gradient was disrupted in *smb-3* root cap. (A) A pie chart illustrating the number of hormone-related differentially expressed genes (DEGs) between Col-0 and *smb-3*. (B) Gene Ontology (GO) term analysis of the DEGs described in Figures1m, highlighting enrichment of auxin signaling-related genes and auxin homeostasis-related genes(B) in the comparison between Col-0 and *smb-3*. (C) Heat map of selected auxin-related DEGs associated with root cap development and formation from the analysis in Figure 2B. (D-G) *DR5rev:GFP* expression pattern in Col-0(D) and *smb-3*(F). Panels E and G show signal intensity profiles measured along the highlighted regions from the quiescent center (QC) to the distal root cap in Col-0(E) and *smb-3*(G). (H-K) *DR5rev:3xVENUS* expression pattern in Col-0(H) and *smb-3*(J). Panels I and K show nucleus intensity heat maps illustrating auxin distribution in Col-0(I) and *smb-3*(K). The heat map were generated using the intensities of each nucleus combined with its cell geometry. (L and M) Expression pattern of DII-VENUS in Col-0(L) and *smb-3*(M). (N-P), Statistical analysis of the fluorescence intensity of *DR5rev:GFP*, *DR5rev:3xVENUS* and DII-VENUS reporters in Col-0 and *smb-3*. Data are presented as mean ± SD. Significant differences were determined using two-tailed Student’s t test (N-P; p<0.0001 is denoted by ****). Scale bars: 50 μm. See also Figure S3

### Inactivation of *SMB* disrupts the auxin gradient in the root cap

It has been demonstrated that auxin gradient within the root cap play critical roles in root cap development and turnover^10^. To test our hypothesis regarding whether SMB is a regulator of auxin-related pathway, we examined auxin distribution in the *smb-3* root cap via auxin reporter lines *DR5rev:GFP*. Levels of auxin in the root cap cells were significantly reduced in *smb-3* in comparison with Col-0 and its concentration gradient was indistinct in the mutant root cap (Figures 2D-2G, and 2N). To further verify the suppressed auxin gradient, we generated *DR5rev:3xVENUS* and DII-VENUS auxin reporter lines in the *smb-3* background. In agreement with the observation in the *smb-3*x*DR5rev:GFP* line, auxin accumulation and its gradient was inhibited in *smb-3xDR5rev:3xVENUS* and *smb-3x*DII-VENUS lines (Figures 2H-2P). These results support the notion that SMB is required for the establishment and maintenance of the auxin gradient in the root cap.

### Perturbation of auxin distribution affects cell differentiation and BLC release in the root cap

Inhibited Golgi remodeling in the *smb-3* root cap could be explained by its stunted auxin gradient as the Golgi restructuring parallels the cell differentiation within the root cap (Figures 1A-1C). We examined BLC release, Golgi structures, and XGA synthesis in the root cap cells after NPA treatment. NPA caused agravitropic growth and interfered with BLC release (Figures 3A-3D, 3H, 3I, and 3K). More than 80% of roots retained more than two layers of BLCs attached to the root tip, compared to only 22% in the control group. Whole-mount immunofluorescence microscopy immunogold labeling with LM8 revealed that NPA-treated root caps accumulated significantly less XGA (Figures 3H and 3I) because Golgi stacks in BLCs did not synthesize XGA (Figures 3F and 1K). The hypertrophied structure and XGA synthesis were restored when the seedlings were removed from the NPA-containing medium (Figures 3G and 1K). When we assessed expression of SMB, miR160a, XGD1 and ALA3 (genes involved in XGA synthesis and swollen vesicle formation) by qRT-PCR, all the genes were downregulated by NPA treatment (Figure 3J).

**Figure 3.**
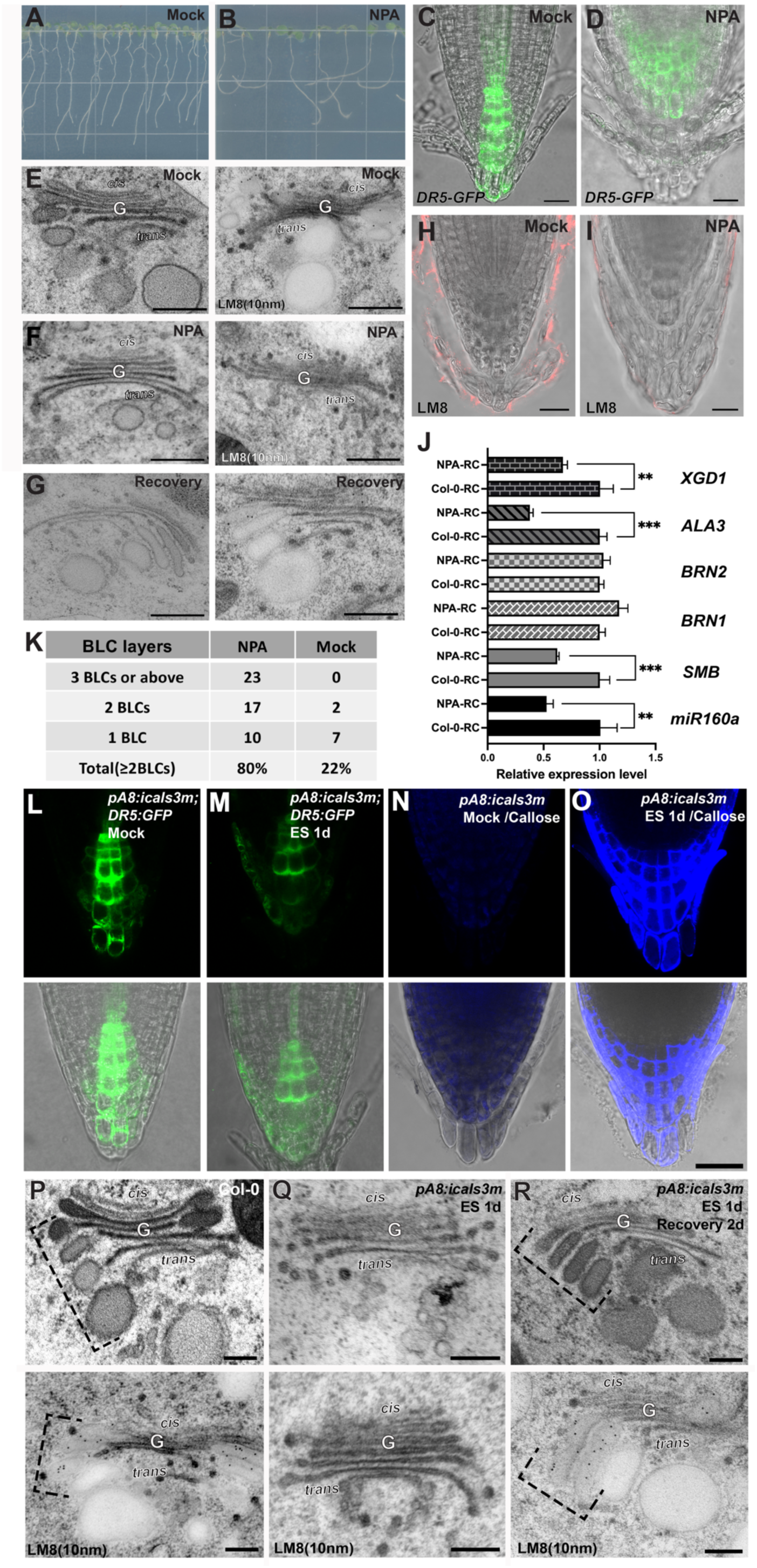
NPA treatment and symplastic communication blockage altered XGA synthesis and Golgi remodeling in root BLCs. (A and B) Root phenotypes of Col-0 grown on 1/2 MS medium with or without 10μM NPA. (C and D) *DR5rev:GFP* expression pattern following 3 days of Mock (C) and NPA(D) treatment . (E-G) Electron micrographs of Golgi stacks (left) and Golgi labeled with an anti-XGA antibody LM8 (right) in BLCs of Col-0 under mock conditions(E), after NPA treatment(F), and recovery after NPA treatment(G). (H and I) Whole-mount immunofluorescence with anti-XGA antibody LM8 under mock (H) and NPA (I) treatment. (J) qRT-PCR analysis of root cap-specific and XGA synthesis genes upon NPA treatment. (K) Statistical analysis of BLC layer number under NPA treatment. (L and M) *DR5rev:GFP* expression in *pA8::icals3m* after 24 hours of 10 μM estradiol induction. (N and O) Aniline blue staining to visualize callose deposition in the root cap of *pA8::icals3m* after 24 hours of 10 μM estradiol induction. (P-R) Electron micrographs of Golgi stacks and Golgi labeled with an anti-XGA antibody LM8 in BLCs of Col-0(P), *pA8::icals3m* after 24 hours of 10 μM estradiol induction (Q) and after 2 days of recovery(R). Data are presented as mean ± SD. Significant differences were determined using two-tailed Student’s t test (J; p<0.05 is denoted by *, p<0.01 by **, p<0.001 by ***). Scale bars: C-D, H-I, and L-O: 50 μm; E-G and P-R: 250 nm. See also Figure S6

In addition to NPA treatment, we employed a callose-inducible system to transiently blocks symplastic communication within the root cap that disrupts auxin gradient formation. Upon estradiol treatment, callose staining of the root cap cell wall significantly increased (Figures 3N and 3O). The enhanced callose deposition attenuated the auxin response in the root cap (Figures 3L and 3M). After 24 hours of estradiol induction, the Golgi swelling in the peripheral cells and BLCs completely disappeared. However, these hypertrophied Golgi stacks reappeared within two days of recovery without estradiol (Figures 3P-3R). Immunogold labeling analysis also showed that XGA-specific gold particles were absent in the estradiol-treated root cap samples, but XGA started accumulating in the Golgi and the cell wall after two days of culture without estradiol, consistent with ultrastructural observations (Figures 3P-3R and 1K). Altogether, these results indicate that the auxin gradient is critical for cell differentiation, Golgi transformation, and BLC turnover in the root cap.

### SMB regulates root cap auxin response by directly binds to the promoter region of auxin signaling-related genes

Our RNA-seq and qRT-PCR analysis results indicated that SMB regulates the expression of auxin signaling-related genes. To validate whether SMB affects the transcription levels of these genes, we performed dual luciferase (LUC) assays with the transient expression system of Arabidopsis PBSD cells. LUC activity dropped in PSBD overexpressing SMB and *ARF10pro:LUC* or *ARF16pro:LUC* reporters (Figures 4E and 4F), suggesting that SMB is involved in the control of *ARF10* and *ARF16* expression. To investigate whether SMB directly interacts with the promoters of these genes, we carried out electrophoretic mobility shift assays (EMSAs) for *ARF10* and *ARF16*, SMB was seen to bind to two SNBE motifs within the promoters of ARF10 (D and G) and ARF16 (A and B) (Figures 4A, 4B, S4A, and S4B). Additionally, SMB was also targeted to the promoter of an ARF10/16 interacting protein, IAA33 (B and D) and an ARF10/16 inhibitor miR160a (B) (Figures 4C, 4D, and S4C). These EMSA results agreed with data from a previous plant cistrome database (Figure S4D)^47^. ARF10 and ARF16 are crucial for root cap maturation and are believed to promote columella cell differentiation^29^. The *arf10arf16* show a severely disorganized root cap morphology due to uncontrolled cell division, as visualized by Edu staining (Figures 4I and 4J). Ultrastructural analysis revealed that most root cap cells in the double mutants retained meristem-like cell architectures including small Golgi stacks with 4-5 cisternae (Figures 4G, 4H, and S5), indicating that ARF10 and ARF16 play roles in columella cell differentiation. These results suggest that SMB modulates the auxin signal outputs by controlling the expression levels of auxin signaling-related genes in the root cap (Figure 4K).

**Figure 4.**
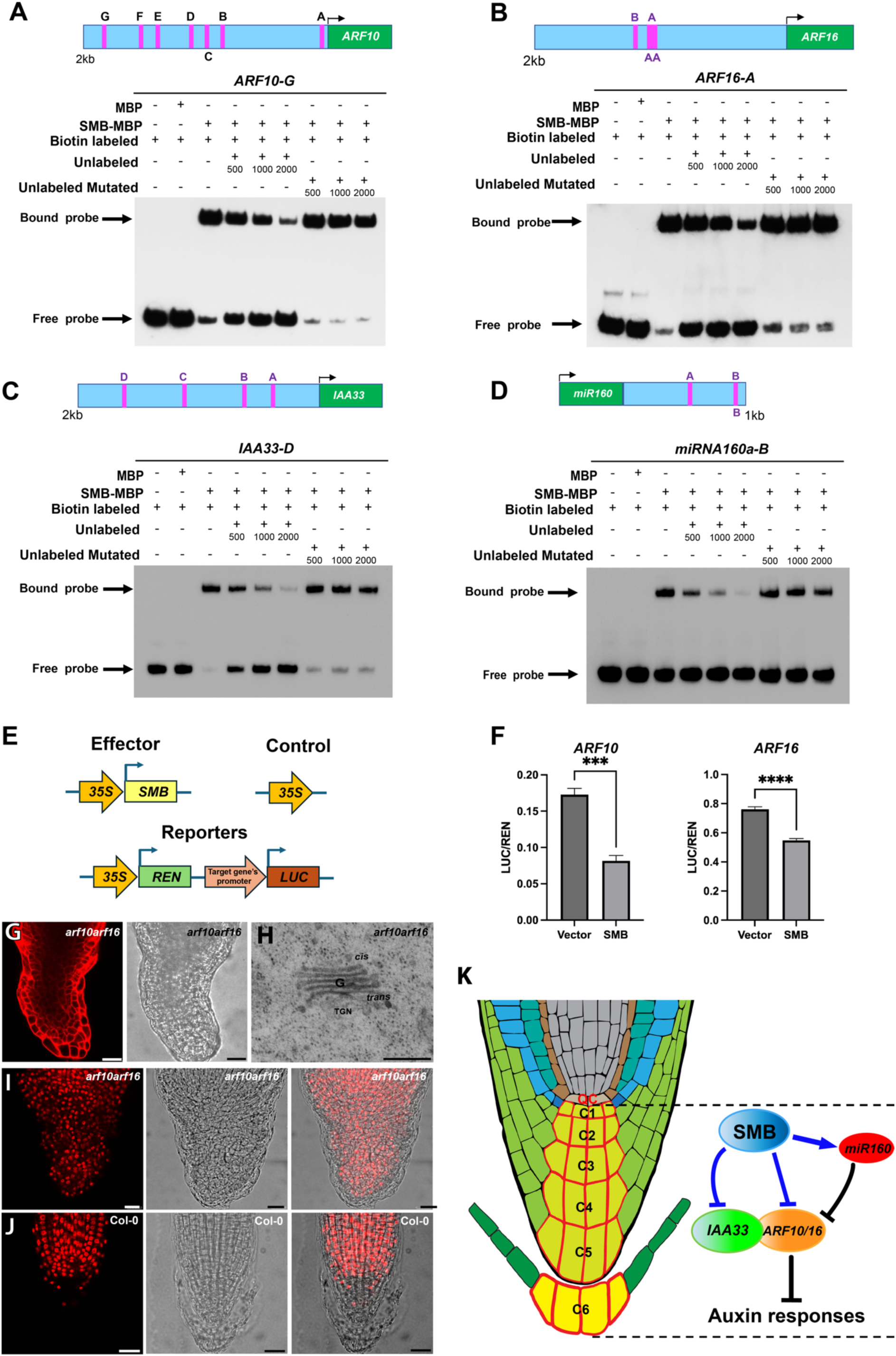
SMB directly binds to the promoter region of auxin signaling-related genes and regulate their transcription. (A-D) EMSA assays showing that SMB binds to Secondary wall NAC Binding Elements (SNBEs) associated with auxin signaling-related genes within 2 kb. Each panel has a diagram in which the transcription starts site (bent arrow) and SNBEs (magenta lines) are marked. SNBEs are labeled with alphabet letters from “A” in the order of their proximity to the transcription start site. SNBEs in the sense and anti-sense directions are indicated by alphabet letters above and below the gene model, respectively. EMSA results of G-SNBE of *ARF10* (a), A-SNBE of *ARF16 (b)*, D-SNBE of *IAA33* (c) and B-SNBE *miRNA-160a* (d) are provided. Unlabeled probes were used in the competition assay (500, 1000, and 2000 fold excess). Arrows indicate the bound probe and free probe. (E) Recombinant constructs for luciferase report assays. The transfection effector, the control, and a reporter plasmids are shown. ∼2kb upstream from the transcription start sites of the target genes (target gene’s promoter) were inserted in front to the firefly luciferase gene (*LUC*). (F) Transient assay for target genes regulated by SMB. Values are means of at least three replicates. Asterisks indicate significant differences between SMB and empty vector. *REN*: *RENILLA* luciferase gene. (G) Disorganized root cap structure of an *arf10arf16* double mutant. A confocal micrograph of the root tip after staining with propidium iodide (left) and its brightfield micrograph (right) are shown. (H) A Golgi stack in an *arf10arf16* root cap cell. (I and J) EdU staining of root cap samples from an *arf10arf16* (I) and Col-0 (J) seedlings. (K) SMB binds to the promoter regions of *ARF10, ARF16, IAA33, and miR160* (blue lines) for auxin signaling within the root cap. Data are presented as mean ± SD. Significant differences were determined using two-tailed Student’s t test (F; p<0.05 is denoted by *, p<0.01 by **, p<0.001 by ***, p<0.0001 by****). Scale bars in G: 50μm; H, 300nm; I-J: 25 μm. See also Figures S4 and S5

### SMB is essential for root cap auxin homeostasis by directly binds to the promoter region of auxin biosynthesis and transport

Root cap growth and cell turnover depend on the local auxin gradient^10^. The hormone gradient in the root cap is maintained by coordinated actions of auxin biosynthesis, transport and inactivation. We uncovered that expression levels of auxin biosynthesis genes (TAA1 and TAR2) and auxin transport genes (PIN3, PIN4 and PIN7) are affected in *smb-3* from our root cap-specific RNA-seq and qRT-PCR analyses (Figures 2 and S3). We conducted dual luciferase assays for the genes in Arabidopsis PBSD as was done for *ARF10* and *ARF16*. SMB increased the transcription levels of *TAR2pro:LUC*, *PIN3pro:LUC*, *PIN4pro:LUC* and *PIN7pro:LUC* (Figure 5E). EMSA for the four genes further showed that SMB binds to their promoter sequences (Figures 5 A-5D) and these results align with the plant cistrome database^47^ (Figure S4D). Interestingly, although *TAA1* transcription was reduced in *smb-3*, SMB failed to bind to the *TAA1* promoter in EMSA tests. This suggests that SMB directly regulates *TAR2* transcription but influences *TAA1* indirectly. Furthermore, both *taa1-1* and *tar2-2* mutants displayed inhibited Golgi remodeling and XGA synthesis (Figure S6).These findings indicated that SMB contributes to the local auxin gradient along the root cap (Figure 5F).

**Figure 5.**
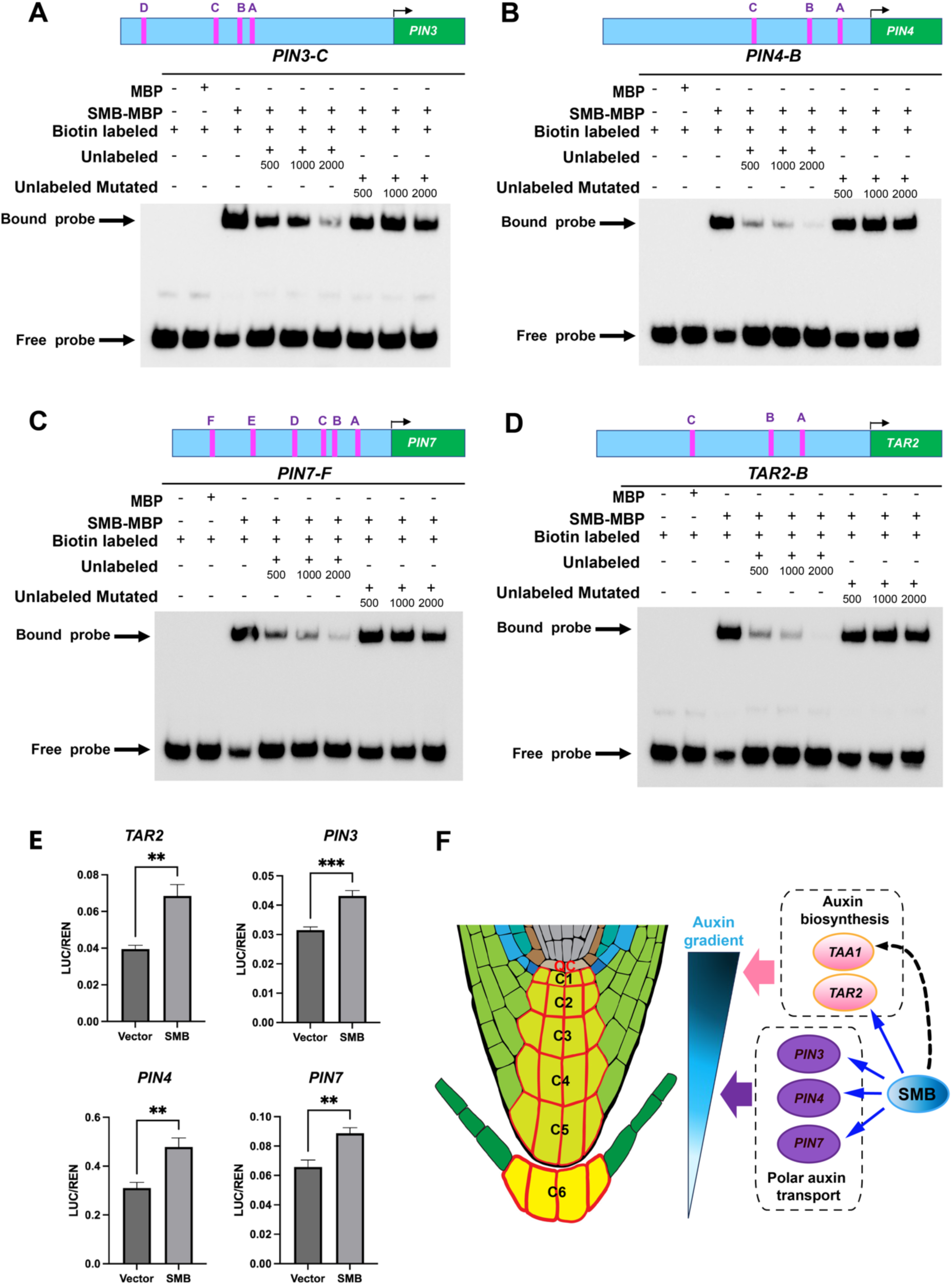
SMB directly binds to the promoter region of auxin biosynthesis and transport genes for their transcriptional regulation. (A-D) EMSA assays showing that SMB binds to SNBEs associated with genes involved in auxin synthesis and polar transport within 2 kb. Each panel has a diagram in which the transcription starts site (bent arrow) and SNBEs (magenta lines) are marked. SNBEs are labeled with alphabet letters from “A” in the order of their proximity to the transcription start site. SNBEs in the sense and anti-sense directions are indicated by alphabet letters above and below the gene model, respectively. EMSA results of C-SNBE of *PIN3* (A), B-SNBE of *PIN4* (B), F-SNBE of *PIN7* (C), and B-SNBE of *TAR2* (D) are provided. Unlabeled probes were used in the competition assay (500, 1000, and 2000 fold excess). Arrows indicate the bound probe and free probe. (E) Luciferase report assay results of the promoter sequences of the genes examined with EMSA assays. (F) SMB regulates expression of auxin biosynthesis and polar transport, including *TAR2, PIN3, PIN4,* and *PIN7* to maintain auxin gradient along the root cap. Data are presented as mean ± SD. Significant differences were determined using two-tailed Student’s t test (E; p<0.05 is denoted by *, p<0.01 by **, p<0.001 by ***). See also Figures S4 and S6

### *smb-3* exhibits reduced sensitivity to NPA

Inhibition of the auxin transports in the root cap by NPA causes agravitropic root growth ^25,26^. Given that transcription of *PIN3, PIN4,* and *PIN7* is impaired in *smb-3*, the NPA’s effect could be altered in *smb-3*. When treated with NPA for 2 days, root tips of *smb-3* exhibited a slight agravitropic response while gravitropic growth was strongly inhibited for Col-0 root tips (Figures 6A-6L, S7 and S9D-S9I). The abnormal growth became more evident on the third day treatment for Col-0 but not for *smb-3* (Figures 6A-6L and S7). The suppressed agravitropic response of *smb-3* was further confirmed in split agar NPA treatment tests (Figure S8). The reduced agravitropic growth of *smb-3* was reversed by expression of *SMB-GFP* via the *SMB* native promoter (Figures 6E, 6F, 6K, and 6L). Interestingly, *smb-3* seedlings exhibited gravitropic response similar to that of Col-0 upon reorienting agar plates (*i.e.* gravti-stimulation), suggesting that NPA-induced abnormal response to gravity is independent of the columella amyloplast-dependent gravitropic growth response (Figures S9A-S9C). The NPA-induced abnormal root growth was mitigated by NAA treatment in both Col-0 and *smb-3* (Figure S10).

**Figure 6.**
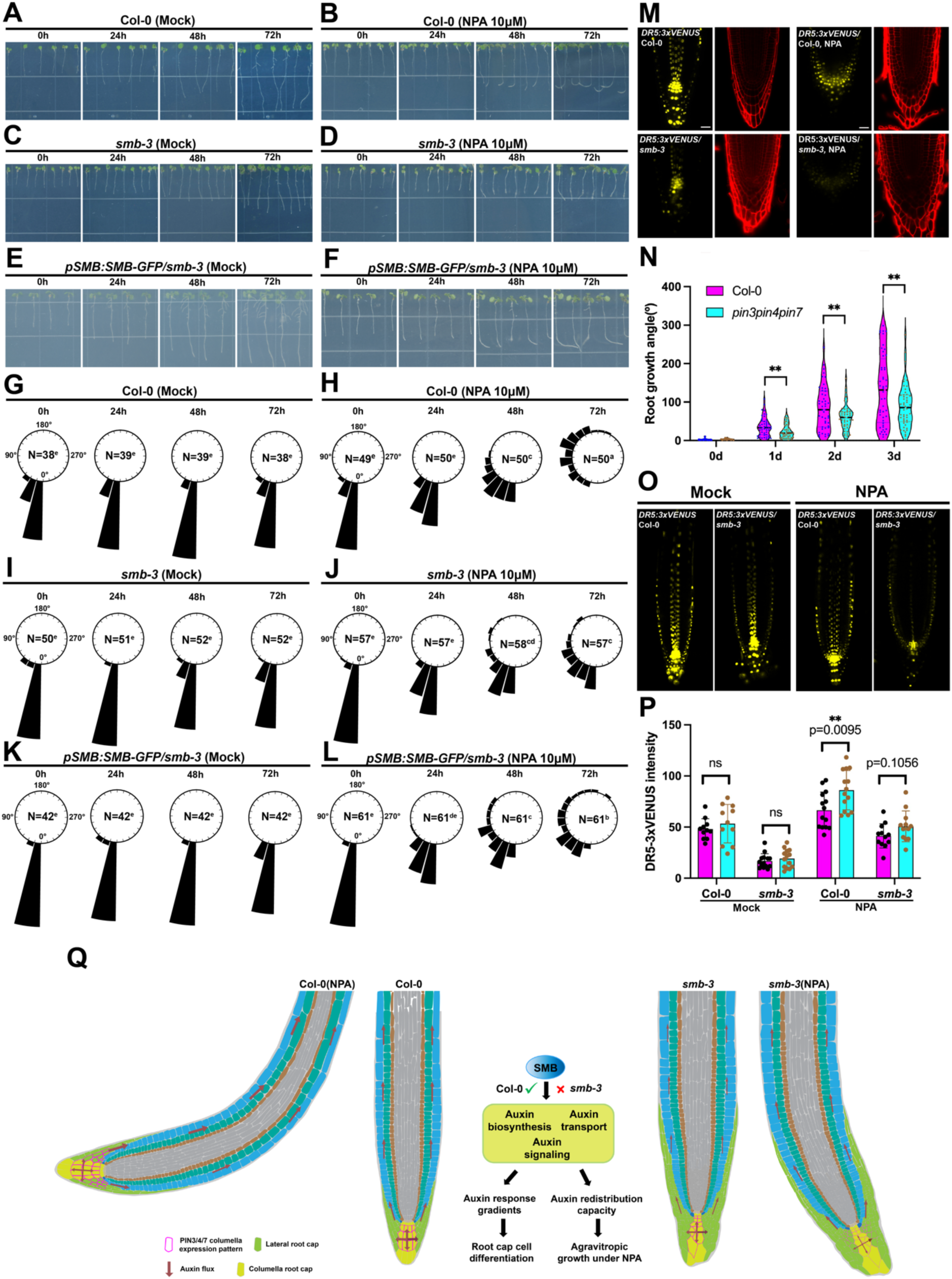
*smb-3* exhibits reduced sensitivity to NPA treatment. (A-F) Time-lapse phenotypic analysis of Col-0(A-B), *smb-3*(C-D) and *pSMB:SMB-GFP/smb-3*(E-F) that grown vertically on 1/2 MS medium with or without 10μM NPA. (G-L) Frequencies of root tip growth angles with or without NPA. The direction of each root tip was measured as the absolute angle relative to the direction of gravity (g). The frequency was calculated as the proportion of root numbers that fell within intervals of 15° relative to the total number of analyzed roots for each line. Bars represent the relative frequency. N indicates the number of independent seedlings, and alphabets denote significant differences (p<0.05, one-way ANOVA by Tukey’s test). (M) Expression pattern of *DR5rev:3xVENUS* in Col-0 and *smb-3* grown vertically on 1/2 MS medium with or without NPA treatment. (N) Violin plot showing *pin3pin4pin7* triple mutant exhibits reduced agravitropic root growth angle under NPA treatment. (O) Confocal images of *DR5rev:3xVENUS* signals in the lateral root cap (LRC) and epidermal cells of Col-0 and smb-3 that grown vertically on 1/2 MS medium with or without 10μM NPA. (P) Statistical analysis of fluorescence intensity of *DR5rev:3xVENUS* in the LRC and epidermal cells on the curvature direction side(cyan) and the other side (magenta) of the root tip under NPA treatment. Under mock conditions, cyan and magenta represent the left and right sides of the root tip, respectively. (Q) A proposed model illustrating SMB functions as a versatile transcription factor that regulates auxin-related pathways to coordinate root cap cell differentiation and control root agravitropic growth under NPA treatment. Data are presented as mean ± SD. Significant differences were determined using two-tailed Student’s t test (N and P; p<0.01 is denoted by **). Scale bars: 25 μm. See also Figures S7-S13

In *smb-3* background, expression of *proPIN3:PIN3-GFP*, *proPIN4:PIN4-GFP* and *proPIN7:PIN7-GFP* was lower than in Col-0 (Figure S11). When treated with NPA, the expression of auxin transporters expanded laterally in the Col-0 but not in *smb-3*, suggesting that auxin reflux to the lateral root cap could be compromised in *smb-3*. This notion was supported by the *DR5rev:3XVENUS* auxin reporter lines that showed that auxin response in the lateral root cap in the Col-0 but not in the *smb-3* backgrounds (Figure 6M). The NPA-induced agravitropic growth was reduced in the *pin3pin4pin7* triple mutant as that in *smb-3* (Figures 6N and S12). These results imply that the inhibited gravity sensing casued by NPA depends on SMB-mediated up-regulation of auxin related genes including *PIN3, PIN4,* and *PIN7*. In agreement with the role of SMB, activation of *SMB* promoted root curvature and the bent angles, amplifying the NPA-induced agavitropic growth (Figure S7).

## DISCUSSION

We have demonstrated that SMB potentiates the auxin gradient and auxin signaling in the root cap, acting as a central regulator of root cap cell differentiation and turnover. Given that SMB functions as a transcriptional activator or as a repressor depending on genes and cell types in the root cap, we propose a model illustrating SMB act as a versatile transcription factor, orchestrating various aspects of the auxin homeostasis and signaling in the root cap(Figures 4K, 5F, and 6Q). Since the auxin gradient directs the timing of root cap cell division, growth, maturation, and detachment, SMB facilitates root growth and redirection in response to the stimuli (Figure 6Q). It has been shown that ethylene stimulates auxin biosynthesis in the roots^48^ while cytokinin antagonizes auxin to regulate the size of the root meristem^49,50^. In agreement with the interactions between hormone pathways in the root tip, many genes involved in hormone-related processes exhibited altered transcriptional activities in *smb-3* (Figure 2A).

### Root cap-specific RNA-Seq revealed significance of SMB in controlling the auxin homeostasis and response in the root cap

It is required to perform spatial RNA-seq analyses to compare expression profiles of genes specifically expressed in specific cell types (Figure 4A). Single-cell RNA-seq provides high-resolution transcriptomic data at the individual cell level. However, the approach has limitations for plant research because it involves cell wall digestion that triggers stress responses in plant cells, complicating results. We developed a root cap-specific RNA-seq method (Figure 1L), identifying over 3,000 DEGs between Col-0 and *smb-3* (Figure 1M). Among these, more than 100 genes were linked to auxin pathways (Figures 2A-2C, and S3). Our method is particularly suited for examining ubiquitously expressed genes in a specific tissue. For example, RNA-seq without root cap dissection did not show significant changes in transcript levels of *PIN3, PIN4,* and *PIN7* between Col-0 and *smb-3* (Figure S4A). By contrast, large drops in their mRNA levels are unambiguous in the dataset after root cap isolation, revealing their activation by SMB in the root cap (Figures S4A and S4B). In addition, Gene expression analysis using the root cap dissection method shows contradictory results for some genes under NPA treatment compared to methods using full-length roots, highlighting the usefulness of this approach for studying specifically regulated genes in the root cap(Figure S3E).

Another noteworthy aspect of our root cap-specific RNA-seq is that a group of genes that were inactive in the Col-0 root cap but became activated in *smb-3*. This finding suggests that SMB may act as a root cap identity gene (Table S2). Root cap-specific genes display distinct ‘Z’ expression patterns when root cap and full-length roots transcripts were compared (Figure S3D; listing well-known root cap-specific genes.) It is possible to discern root cap-specific genes or genes that are broadly expressed but show significant enrichment in the root cap by comparing these two datasets (Table S1). We believe that our tissue-specific transcriptomics method offers a complementary approach to general RNA-seq and single-cell RNA-seq (SC RNA-seq) for advancing our understanding of root cap cell development and renewal.

### Root cap auxin gradient and cell differentiation in the central root cap

Root cap serves as an excellent model for studying the relationship between auxin gradient and organ development in plants^51^. The central root cap of Arabidopsis consists of four or five tiers of columella cells with BLCs at the surface, making it an ideal system for study cell differentiation gradient, and programmed cell death (PCD). Auxin forms a maximum at stem cell niche (SCN), which helps maintain its identity, and set up an auxin gradient from the SCN to the distal root cap, parallelling the cell differentiation. We hypothesize that the auxin gradient drives the maturation of columella initial cells, columella cells, and eventually into BLCs that will be shed and undergo PCD. In each cell type, Golgi stacks exhibit unique ultrastructural features, serving as the cell identity marker in the central root cap (Figure 1).

It has been reported that the function of SMB is determined by its expressive level, with strong *35S:SMB-GR* activation leading to dramatic growth inhibition^35^. When we switched on 35S:SMB-GR with varying concentrations of DEX (Figures S2B-S2D), we observed that low DEX levels increased the degree of Golgi swelling and amounts of XGA. However, strong activation of *35S:SMB-GR* severely disrupted Golgi morphology and XGA synthesis (Figures S2B-S2E), in agreement with the previous study. Given the role of SMB in root cap auxin-related pathway. Strong activation of SMB may also severely disrupt auxin gradient formation, leading to dramatic growth inhibition. These results indicate that the Golgi-mediated secretion in the root cap is tightly controlled, auxin and appropriate SMB expression levels playing crucial roles in the process. Exploring the regulatory network that maintains appropriate SMB expression levels will be an interesting focus for future research.

Plasmodesmata (PD)-dependent symplastic transport regulates a wide range of developmental processes in plants^52–54^. Researchers have utilized an inducible module (icalsm3) driven by a root cap-specific promoter to block symplastic communication by synthesizing callose in PD^55^. Callose accumulation in the PD of root cap cells disrupted auxin transport and gradient formation, which in turn severely impaired root development in Arabidopsis^55^. The DEX-induced PD clogging in the root cap inhibited XGA synthesis more severely than auxin transport inhibitors or inactivation of SMB (Figures 1K and 3Q). This suggests that PD-dependent intercellular transport mediates cell-cell signalling for the root cap cell differentiation beyond auxin transport.

### SMB contributes to the agravitropic response under NPA treatment

NPA specifically binds to auxin transporter proteins. The chemical may mimic or interfere with endogenous molecules that plant cells produce for controlling auxin transport^23,24^. Our fundings suggest that SMB may function as a regulator of the endogenous counterpart of NPA, as the SMB loss-of-function mutant shows decreased sensitivity to NPA treatment. Additionally, we observed that Col-0 seedlings exhibit varying sensitivity to different concentrations of NPA, with the highest sensitivity observed at 5 μM NPA (Figure S13). The root cap functions as a local auxin sink^56^. The larger bending angle observed at low NPA concentrations(5 μM) may be due to partial disruption of auxin transport, which enhances the asymmetric auxin distribution between the two lateral root regions. However, at high NPA concentrations, auxin transport is likely be blocked completely, leading to a reduction in asymmetric auxin distribution and thereby bending. In the *smb-3*, by contrast, auxin biosynthesis, transport and signaling are compromised, significantly impairing the ability to concentrate auxin in one of the two lateral root region. This explains why the *smb-3* mutant exhibits weaker gravitropic response(Figures 6 and S13).

### Limitations of the study

In this study, we utilized a root cap-specific sample preparation method to demonstrate that the root cap-specific transcription factor SOMBRERO (SMB) serves as a central regulator of multiple auxin pathways critical for the establishment and maintenance of the auxin gradient. This gradient is essential for root cap development and its responses to external stimuli. SMB functions as a versatile regulator, orchestrating the expression of genes across various auxin pathways. However, the precise mechanisms by how SMB coordinates these gene expression programs to sustain the auxin gradient remain unclear. Additionally, while root cap development is tightly regulated, the potential interactions between SMB and other root cap regulators in controlling the timing of root cap cell development, renewal and responses to external stimuli require further investigation.

## RESOURCE AVAILABILITY

### Lead contact

Further information and requests for resources and reagents should be directed to and will be fulfilled by the lead contact, Byung-Ho Kang

### Materials availability

All plant materials generated in this study are available from the lead contact with a completed materials transfer agreement.

### Data and code availability

All data associated with this study are present in the paper or the supplemental information. This paper does not report original code.

## ACKNOWLEDGEMENTS

This work was financially supported by grants from the Research Grant Council of Hong Kong (GRF14113921, GRF14109222, GRF14110823, GRF14113424, N_CUHK462/22, and C4014-23G) awarded to B.-H.K. We thank Professor Tom Beeckman from the University of Ghent, Professor Wu Shuang from Fujian Agricultural and Forestry University, and Professor Xiang Fengning from Shandong University for providing transgenic lines and mutants described in the methods of this paper. We also thank Professor Chen Haodong from Tsinghua University for sharing the script used to generate circular histograms.

## AUTHOR CONTRIBUTIONS

P.W., B.-H.K, and Z.L. designed research; P.W., Z.L. and Y.Z. performed the experiments; P.W., Z.L., Y.Z., J.H. and B.-H.K. analyzed data; P.W. and B.-H.K. wrote the paper.

## STAR METHODS

### Experimental model and study participant details Plant materials and growth conditions

All *Arabidopsis thaliana* (L.) mutant and transgenic lines were created in the wild-type Col-0 background unless otherwise noted. The *smb-3* (N657070), *DR5rev:GFP*(N9361), *pPIN4:PIN4-GFP*(N9576), *DR5rev:3xVENUS*(N799364), *taa1-1*(N66987), t*aa1-1xDR5rev-GFP*(N66989) and *tar2-2*(N16450) were obtained from Arabidopsis Biological Resource Center (ABRC, Columbus, OH, USA). The *arf10arf16* mutant was provided by Dr. Xiang Fengning^29^. Dr. Wu shuang provided the *pA8:icals3m*, *pA8:icals3m* x *DR5rev-GFP*, *pPIN3:PIN3-GFP*, *pPIN7:PIN7-GFP* lines. The *35S:SMB-GR* line was obtained from Dr. Tom Beeckman^17^. The *smb-3xDR5rev-GFP*, *smb-3xDR5rev-3xVENUS* and *smb-3*xDII-VENUS was generated by crossing *smb-3* with *DR5rev-GFP*, *DR5rev-3xVENUS,* and DII-VENUS, respectively. pSMB:SMB-GFP/smb-3 line was created by transforming the *pSMB:SMB-GFP* vector into the *smb-3* mutant. The *pin3pin4pin7* triple mutant was described previously^57^. Seedlings were grown vertically on half-strength Murashige and Skoog (1/2 MS) medium containing 0.8% agar without sucrose, at 22 °C under the long-day condition (16-hour light/8-hour dark cycle).

### Method details

#### Root cap-specific RNA extraction, RNA-seq and RT-qPCR

Root tips (0.15–0.25 mm) from 7-day-old seedlings were dissected on a small planchette under a stereo microscope. The root tips were quickly cut, and the planchette with the root caps was immediately transferred to an Eppendorf tube containing liquid nitrogen (LN2). After collecting several hundred root caps, ∼20 pre-cooled steel balls were added to the tube, and the mixture was ground using a vortex mixer. Total root-cap-specific RNA was then extracted using the RNeasy Plant Mini Kit (Qiagen, Cat. 74904).

For RNA sequencing (RNA-seq), library construction, quality control, and RNA sequencing were conducted by BGISEQ platform. Each sequencing run yielded between 48 to 53 million raw reads per sample. The sequencing read counts were quality-checked and trimmed to remove adaptor contamination and low-quality reads. Cleaned reads were aligned to the Arabidopsis reference genome (TAIR10), and gene read counts were generated using HTSeq-count with default parameters. Differential gene expression analysis was performed using the DESeq2 package, identifying differentially expressed genes (DEGs) based on a p-value < 0.05 and a fold change > 1.414. Gene ontology annotation of DEGs was carried out using Dr. TOM, and volcano and bubble plots were generated using the same software.

For RT-qPCR, cDNA was synthesized from ∼1 μg of total root-cap-specific RNA using the Bio-Rad iScript™ gDNA Clear cDNA Synthesis Kit (Bio-Rad, 1725035). SYBR Green PCR Master Mix (Bio-Rad,1725271) was employed for the RT-qPCR reaction, which was performed on a CFX96™ Real-Time PCR Detection System (Bio-Rad). NADPH was used as a reference gene, with at least three biological replicates for each sample. All RT-qPCR primers are listed in Table S3.

#### Electrophoresis mobility shift assay (EMSA)

For recombinant protein production, the *SMB* gene was first cloned into *pMal-C2X* vector in frame with a C-terminal MBP tag. The MBP-SMB fusion protein was then purified from *Escherichia coli* BL21 cell line using amylose agarose beads. Oligo probes were synthesized and labeled with biotin using Biotin 3’ End DNA Labeling Kit (Thermo Scientific, Cat. 89817). The labeled single strand probes were annealed into double strands and EMSA was performed using LightShift^®^ Chemiluminescent EMSA Kit (Thermo Scientific, Cat. 20148). For the binding reaction, 2 μg of purified MBP-SMB was incubated with biotin-labeled promoter probes at 25°C for 40 min according to the manufacturer’s instructions. To assess binding specificity, 500-, 1000-, and 2000-fold excess of unlabeled (cold) probes (without 3’-biotin labeling) and mutated cold probes of the SNBE consensus sequence were added to the reaction mixtures for competition assay with the biotin-labeled probes. The binding efficiency was measured with Chemiluminescent Nucleic Acid Detection Module Kit (Thermo Scientific, Cat. 89880). All DNA probes synthesized for EMSA are listed in Table S3.

#### Transient expression

The protein coding sequences of *SMB-GFP* and *GFP* were cloned into the pBI221 vector under the control of the CaMV35S promoter to serve as effector constructs. Promoter sequences of *ARF10*, *ARF16*, *PIN3*, *PIN4*, *PIN7*, *TAR2*, and *TAA1* were amplified and inserted into the pGreenII 0800-LUC reporter vector. Both recombinant plasmids were then transferred into Arabidopsis *PSBD*. Firefly luciferase activity (LUC) and Renilla luciferase activity (REN) were measured using the Dual-Luciferase Reporter Assay System (Promega, Cat. E1910). The ratios of LUC to REN were calculated for determining final transcriptional activities. Experiments were performed in triplicate, and significant differences were evaluated using Student’s *t*-test.

#### Gravitropic phenotype analyses

For agravitropic phenotype analyses, seedlings were grown vertically on 1/2 MS plates for 4 days, then transferred to 1/2 MS plates supplied with or without NPA and grown vertically for an additional three days. The seedings were scanned every 24h using a photo scanner (Epson Perfection V800 Photo scanner). Root tip curvature was measured as the angle between the direction of primary root tip and gravity vector. For the gravity response analyses, seedings were grown vertically on 1/2 MS plates for 4 days, transferred to 1/2 MS plates supplied with or without NPA and then rotated by 90 degrees and grown vertically for an additional day. The root tip angels were measured as the angles between the directions of primary root tip and either the horizontal or gravity vector, respectively. Root growth angles were measured using ImageJ software. The distribution frequency of root curvatures was plotted by the R statistics program(http://www.rproject.org).

#### NPA containing Split-agar assay

The split-agar medium was prepared according to a previously described method(Galvan-Ampudia et al., 2013). A square petri dish was first filled with 1/2 MS medium containing 0.8% agar. Once solidified, the bottom right half of the medium was removed and replaced with 1/2 MS medium supplemented with 0.8% agar and 10 μM NPA. Four-day-old seedlings were then transferred to the split-agar medium plate and grown vertically for an additional days.

#### High-pressure freezing, freeze substitution and resin embedding

High-pressure freezing, freeze substitution, resin embedding were performed as previously described in Kang (2010). Briefly, the root tips were dissected and cryofixed with a EM ICE (Leica Microsystems). After high-pressure freezing, samples were freeze-substituted at -80 °C for 24h in an EM AFS2 freeze-substitution device (Leica Microsystems). For ultrastructural analysis, samples were processed in anhydrous acetone with 2% osmium tetroxide (OSO_4_). Following freeze substitution and multiple rinses, samples were embedded in Epon resin and polymerized at 60 °C. For immunogold labeling or immunofluorescence, root tips were prepared in anhydrous acetone containing 0.25% glutaraldehyde and 0.1% uranyl acetate. The samples were then gradually warmed to -45 °C, held at this temperature, and embedded in Lowicryl HM20 before being cured under ultraviolet (UV) light.

#### Immunogold labeling, Immunofluorescence and CLEM

Ultrathin sections (90–150nm) of HM20-embedded samples were mounted on formvar-coated nickel slot grids. First, grids were placed on drops of 0.1 N HCl for 10 minutes, with the section side in contact with the solution. The grids were then transferred to 2% non-fat milk dissolved in phosphate-buffered saline (PBS) with 0.2% Tween-20 (PBST) and incubated for 30 minutes for blocking. Following this, sections were incubated with primary antibodies. After primary antibody incubation, sections were thoroughly washed in containing 1% non-fat milk in PBST. Next, sections were incubated with secondary antibodies diluted 1:10 in PBST containing 0.5%. These secondary antibodies were conjugated to gold particles of 15 nm, 10 nm, or 6 nm, following the method described by Kang (2010). For immunofluorescence and CLEM, PBS was used as the buffer solution, and grids were kept in the dark before confocal or microscopy imaging.

#### Transmission electron microscopy, electron tomography and 3D model reconstruction

For general transmission electron microscopy (TEM) analysis, ultrathin sections (90– 100 nm) were prepared for ultrastructural examination and analyzed using a Hitachi H-7650 at 80 kV. For electron tomography, thicker sections (250–300 nm) were collected and double-coated with formvar. Tilt series were captured at magnifications of 14,500x or 13,000x, ranging from +60° to -60° with 1.5° increments using an FEI F20 TEM. Tomograms were reconstructed using the etomo interface of the IMOD software package, and calculations for the tomogram and 3D model reconstruction were performed with the 3Dmod graphic module of the same software package.

#### Estradiol induction and callose staining analysis

Estradiol was dissolved in DMSO to prepare 1/2 MS plates containing β-estradiol at a final working concentration of 10 μM. Seedlings were grown vertically on estradiol-free plates for 5 days, then transferred to estradiol-containing plates for an additional 3 days prior to electron microscopy (EM) and confocal laser scanning microscopy imaging. For recovery analysis, after 3 days of growth on the estradiol-containing plates, seedlings were transferred back to estradiol-free plates for another 3 days of recovery.

Callose staining was performed as previously described (Ingram et al., 2011; Li et al., 2022). In brief, seedlings were stained in 67 mM K₃PO₄ and 1% aniline blue for 20 minutes in the dark, and then imaged using a Leica SP8 confocal microscope.

#### Whole-mount immunofluorence

Whole-mount immunofluorescence was conducted following established protocols (Sauer et al., 2006). Seven-day-old seedlings were fixed with 4% paraformaldehyde at room temperature. Following fixation, tissue permeabilization and blocking were carried out before probing with LM8 (Kerafast, ELD012) at 4 °C overnight. After thorough washing with PBS, the samples were incubated with a secondary antibody (Invitrogen; A-11077) at 37 °C for 3 hours for fluorescence microscopy imaging.

#### Edu staining

EdU staining was conducted as previously described (Hong et al., 2015). Briefly, after 4 days of germination, seedlings were transferred to 1/2 MS medium with 0.8% agar supplemented with 10 µM EdU and allowed to grow vertically for 24 hours to incorporate EdU into newly synthesized DNA. Following incubation, samples were fixed in 4% paraformaldehyde and a Click-iT reaction cocktail (Invitrogen Click-iT EdU Imaging Kit, Cat. C10340) containing fluorescent azide was then applied to visualize EdU-labeled DNA. The stained seedlings were thoroughly washed and imaged using a fluorescence microscope.

### Quantification and statistical analysis

The fluorescence intensity and root growth angle were quantified or measured with ImageJ. Statistical analysis were performed in GraphPad Prism 10(GraphPad Software, La Jolla California USA). Statistical details of experiments were indicated in the corresponding figure legends.

## SUPPLEMENTAL INFORMATION

Document S1. Figures S1-S13

Table S1. Potential root cap specific genes or root cap enriched genes

Table S2. GENES not expressed in the Col-0 root cap but highly expressed when inactive SMB.

Table S3. Primers and probes used in this paper.

---

**Figure S1.**
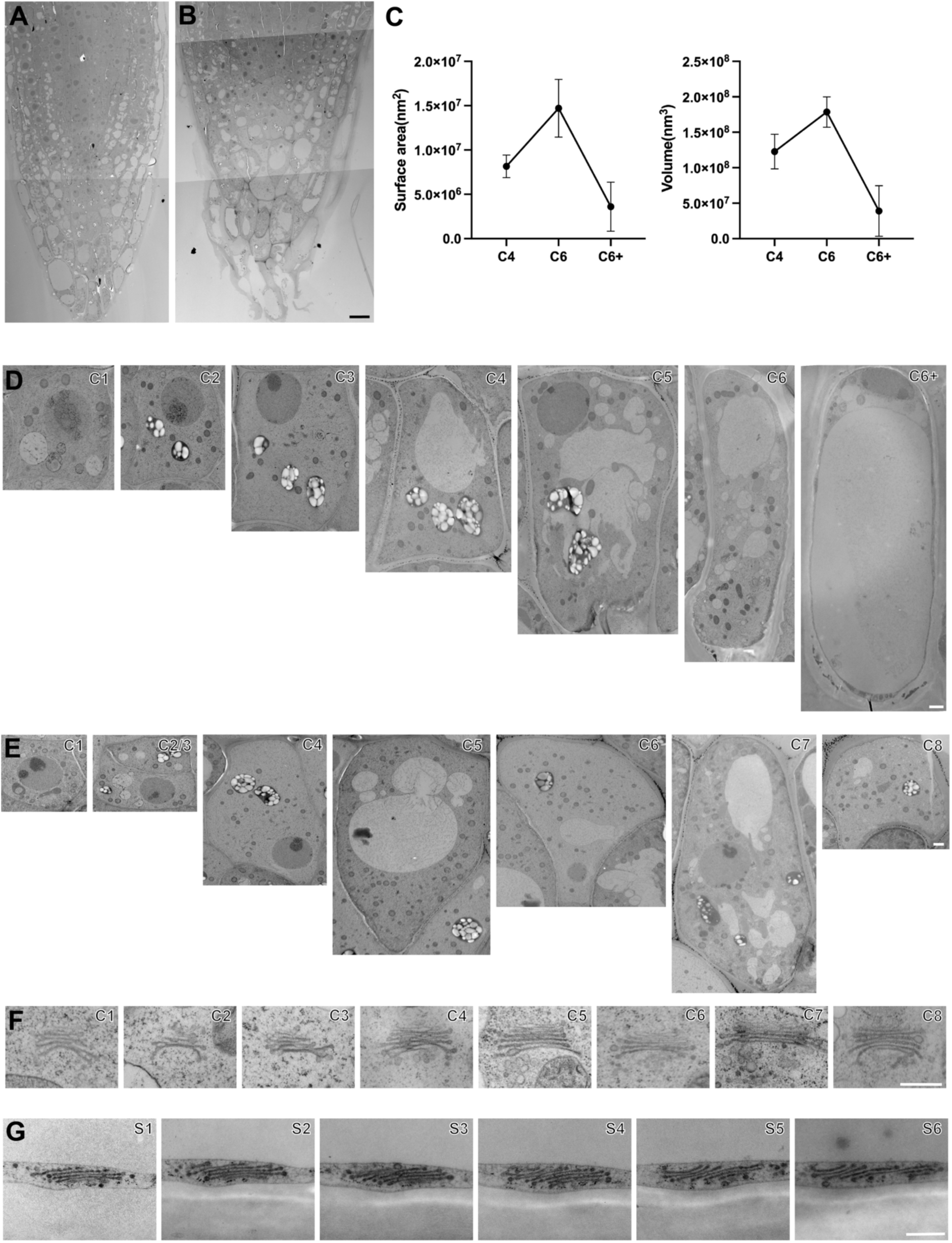
Transmission electron micrographs of *Arabidopsis* root tip. (A and B) Low-magnification electron micrograph (montage) of a longitudinal section of an *Arabidopsis* root tip of Col-0(A) and *smb-3*(B). (C) Volume and surface area analysis of Golgi from late columella cells to highly vacuolated border-like cells (BLCs). (D) Electron micrographs of columella stem cells(C1), columella cells(C2-C5), BLCs(C6) and highly vacuolated BLCs(C6+) of Col-0. (E) Electron micrographs of columella stem cells to outermost BLCs(C1-C8) of *smb-3*. (F) Electron micrographs of Golgi stacks in the root columella cells of smb-3 root cap shown in panel E. (G) Electron micrographs of Golgi stacks obtained from six serial sections in highly vacuolated BLCs of Col-0. Data are presented as mean ± SD. Scale bars in A-B: 10 μm; D-E: 1 μm; F-G: 400nm.

**Figure S2.**
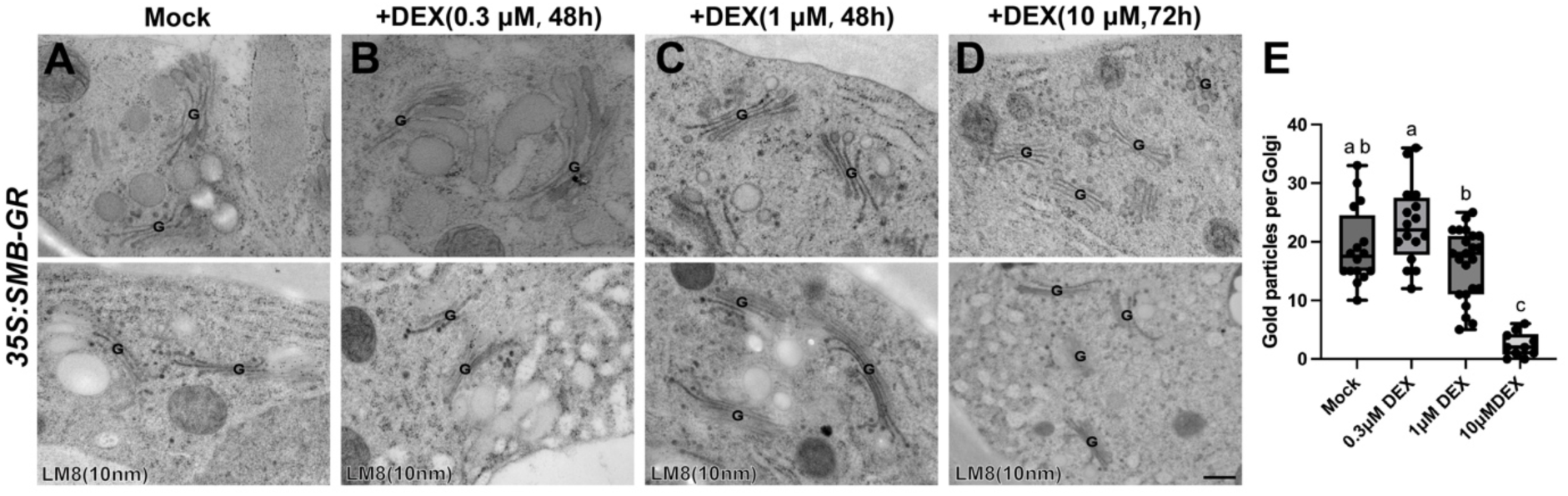
Ultrastructural analysis of Golgi stacks of *35S:SMB-GR* root BLCs under DEX treatment. (A-D) TEM images showing Golgi stacks(upper) and Golgi labelled with an anti-XGA antibody LM8(lower) under varying concentrations of DEX treatment. (E) Statistical analysis of XGA-specific gold particles under different concentrations of DEX treatment. Scale bars in A-D: 400nm. Data are presented as mean ± SD. Alphabets indicate significant differences (p<0.05, one-way ANOVA with Tukey’s test).

**Figure S3.**
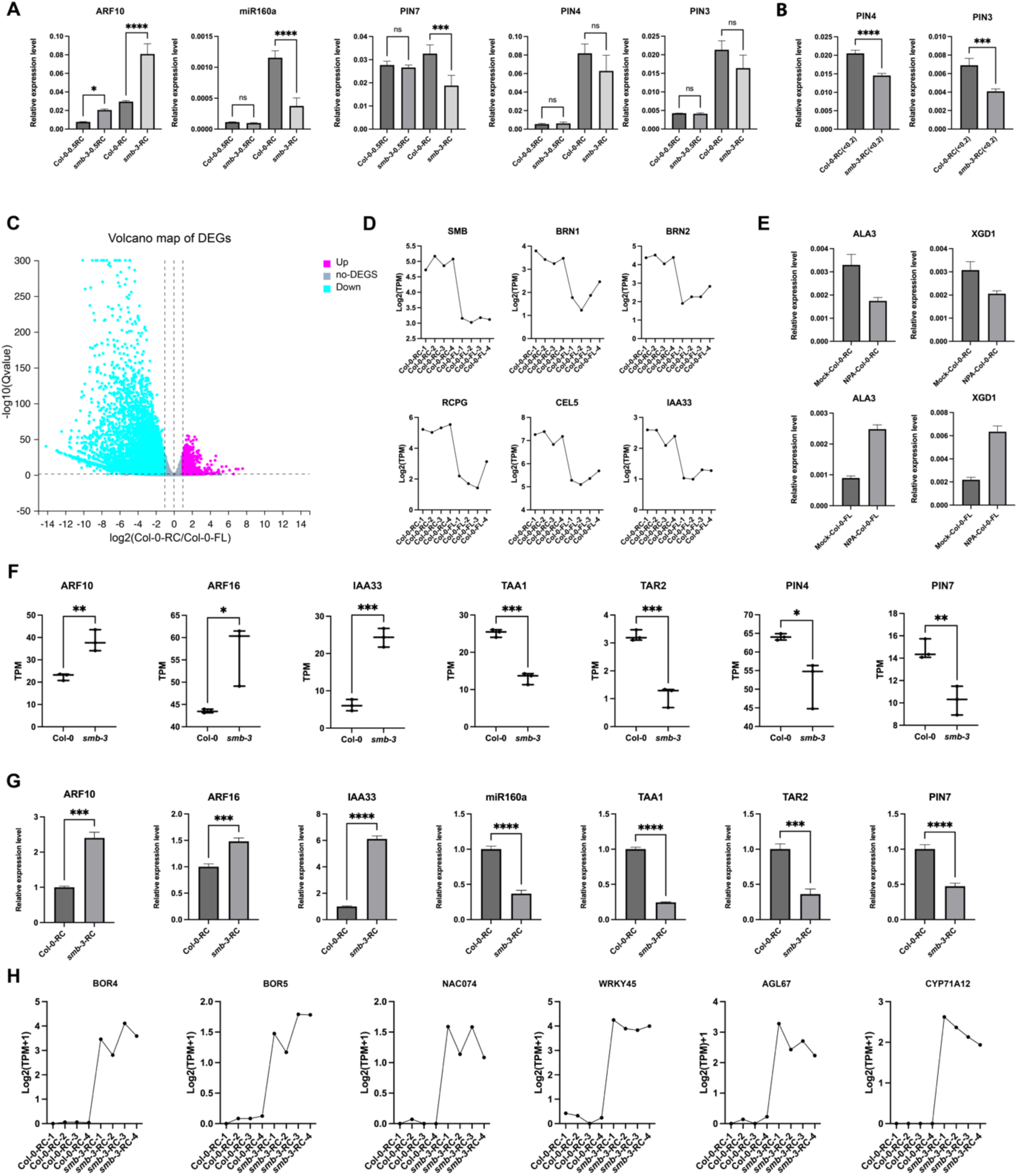
A root cap-specific RNA-seq reveal genes that specifically regulated by root cap TF SMB. (A and B) Comparison of gene expression in different length of root cap by qRT-PCR analysis. “RC” represents root caps ranging from 0.15-0.25mm, “0.5RC” corresponds to root caps approximately 0.5cm in size, and “RC(<0.2)” indicates root caps predominantly smaller than 0.2 mm. (C) Volcano plot of differentially expressed genes (DEGs) between Col-0(RC) and Col-0(full-length root). (D) Several well-known root cap specific genes recognized from up-regulated DEGs in volcano map of panel C. (E) Comparison of gene expression pattern between NPA treated root cap and full-length root. (F) Representative candidate SMB-regulated DEGs identified from RNA-seq data. (G) qRT-PCR analysis validating the expression patterns of candidate genes identified from RNA-seq results. (H) Representative genes that are not expressed or are scarcely expressed in the root cap of Col-0 but are highly expressed in the root cap of *smb-3*. In panels A, B, F and G, values are presented as means ± SD; Statistical significance was assessed using a two-tailed Student’s t test (p<0.05 is denoted by *, p<0.01 by **, p<0.001 by ***, p<0.0001 by****).

**Figure S4.**
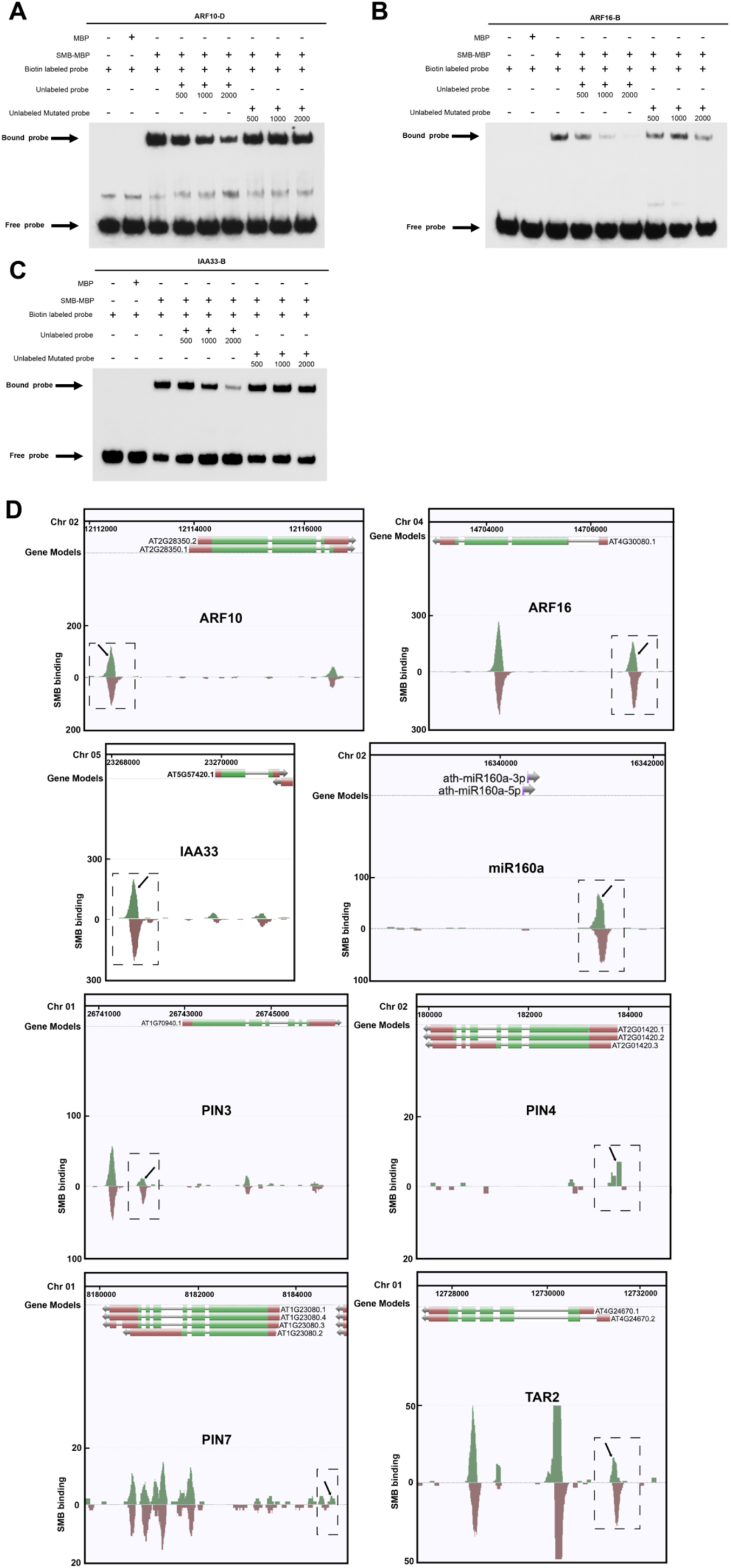
SMB directly binds to the promoter region of auxin-related genes. **(A-C)** EMSA assays showing that SMB binds to the SNBEs in the promoter region of ARF10, ARF16, and IAA33. EMSA results of D-SNBE of *ARF10* (A), B-SNBE of *ARF16* (B), and B-SNBE of *IAA33* (C) are provided. The MBP-labeled protein and biotin labelled probes are indicated in **(A-C)**. Unlabeled probes were used in the competition assay(500, 1000, and 2000 fold excess). Arrows indicate the bound probe and free probe. (**D**) The potential SMB binding sites of auxin-related genes based on plant cistrome database^32^

**Figure S5.**
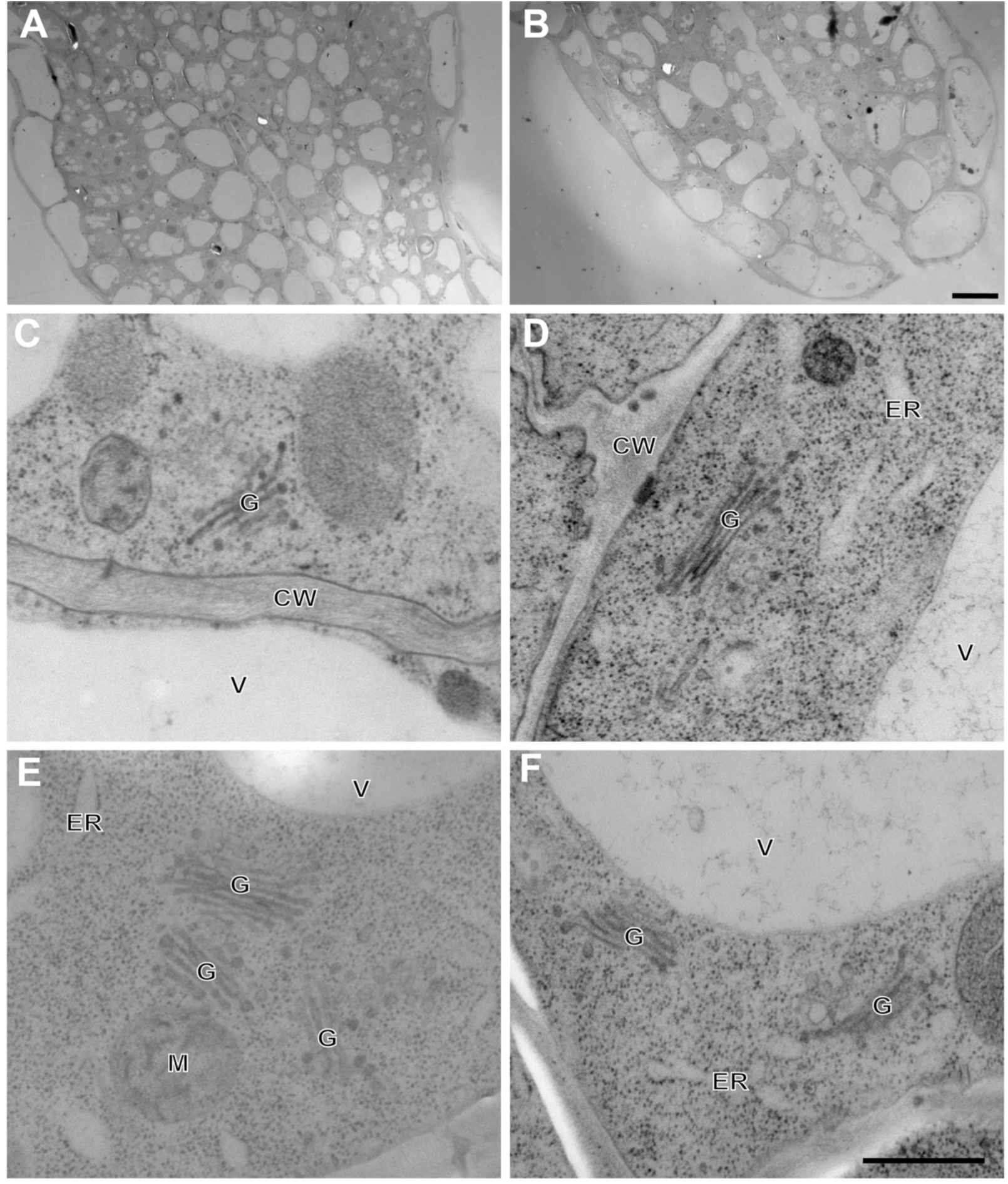
Transmission electron micrographs of *arf10arf16* root cap cells. (A and B) Low-magnification electron micrographs of *arf10arf16* root cap. (C-F) Electron micrographs of Golgi stacks in the root cap of *arf10arf16*. Scale bars in A and B: 10 μm; C-F: 500nm.

**Figure S6.**
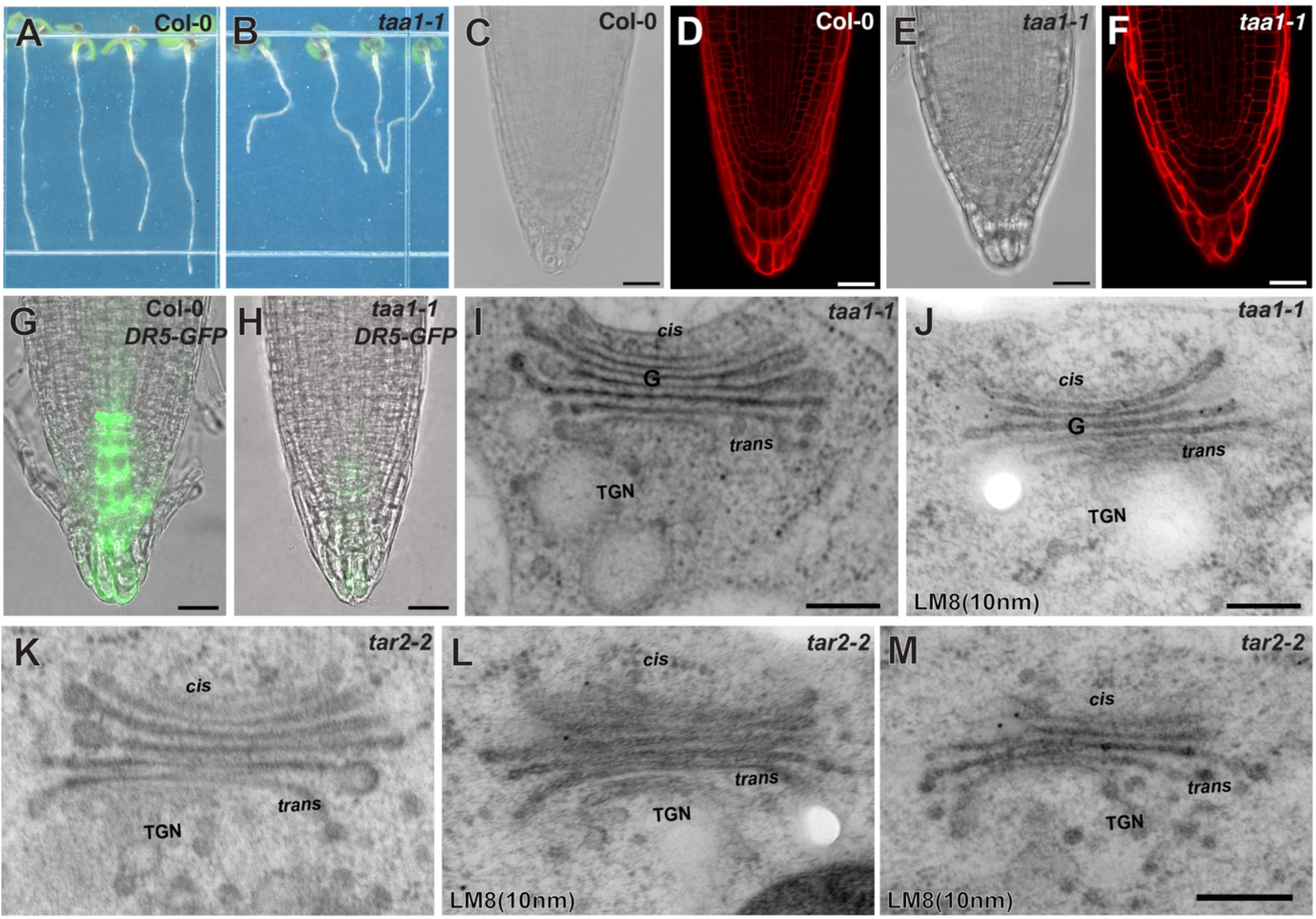
Light and electron microscopy imaging of TAA1 and TAR2 loss of function mutant. (A and B) Light microscopy image of Col-0 and *taa1-1* seedlings grown vertically on 1/2 MS medium. (C-F) Bright field and PI-stained root tip of Col-0 and *taa1-1.* g-h, *DR5rev:GFP* expression pattern in Col-0(G) and *taa1-1*(H). (I and J) Electron micrographs of Golgi stacks(I) and Golgi labeled with XGA antibody LM8(J) in BLCs of *taa1-1*. (K-M) Electron micrographs of Golgi stacks(K) and Golgi labeled with XGA antibody LM8 in BLCs of *tar2-2*. Scale bars in C-H: 50 μm and G-M: 250 nm.

**Figure S7.**
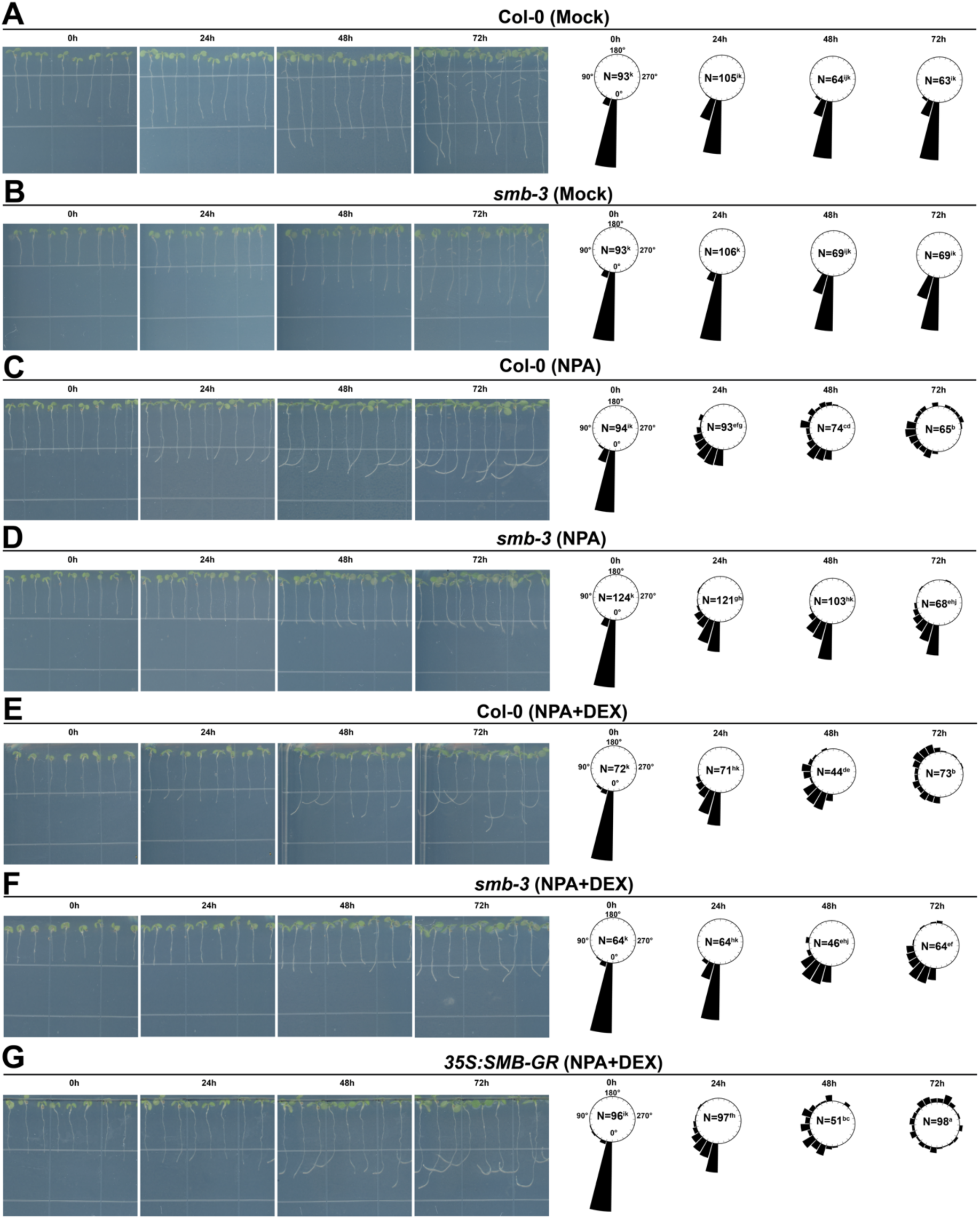
SMB is required for the root agravitropic growth upon NPA treatment. (A-G) Time-lapse imaging of Col-0, *smb-3* and *35S:SMB-GR* that grown vertically on 1/2 MS medium, with or without 10μM NPA and 1μM DEX. Left panel, representative images of seedlings. Right panel, Frequencies of root tip growth angles. The bending angle of each root tip was measured as the absolute value relative to the direction of gravity (g). The frequency was calculated as the proportion of roots falling within 15° intervals relative to the total number of analyzed roots for each line. The bar represents relative frequency. N indicates the number of independent seedlings used in the experiment, and letters indicate significant differences (p<0.05, one-way ANOVA with Tukey’s test).

**Figure S8.**
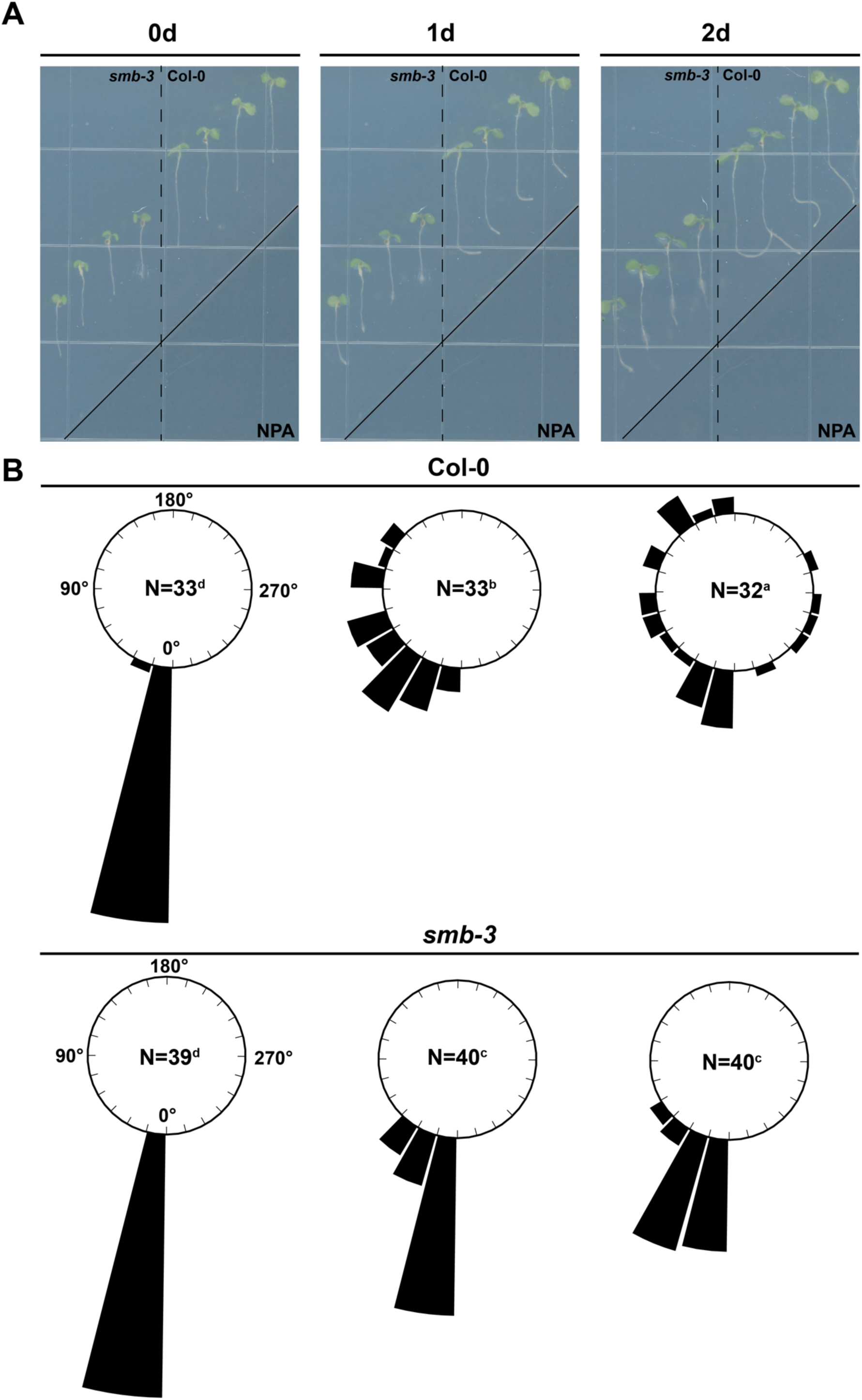
SMB is required for root agravitropic growth in response to NPA treatment. (A) Time-lapse imaging of Col-0, *smb-3* seedlings that transferred to split-agar medium. The upper left side contains 1/2 MS medium, while the right bottom side contains 1/2 MS medium with NPA. The solid line indicates the boundary between 1/2 MS medium and 1/2 MS medium with NPA. (B) Frequencies of root tip bending angles in split-agar system. The bending angle of each root tip was measured as the absolute value relative to the direction of gravity (g). The frequency was calculated as the proportion of roots falling within 15° intervals relative to the total number of analyzed roots for each line. The bar represents relative frequency. N indicates the number of independent seedlings used in the experiment, and letters indicate significant differences (p<0.05, one-way ANOVA with Tukey’s test).

**Figure S9.**
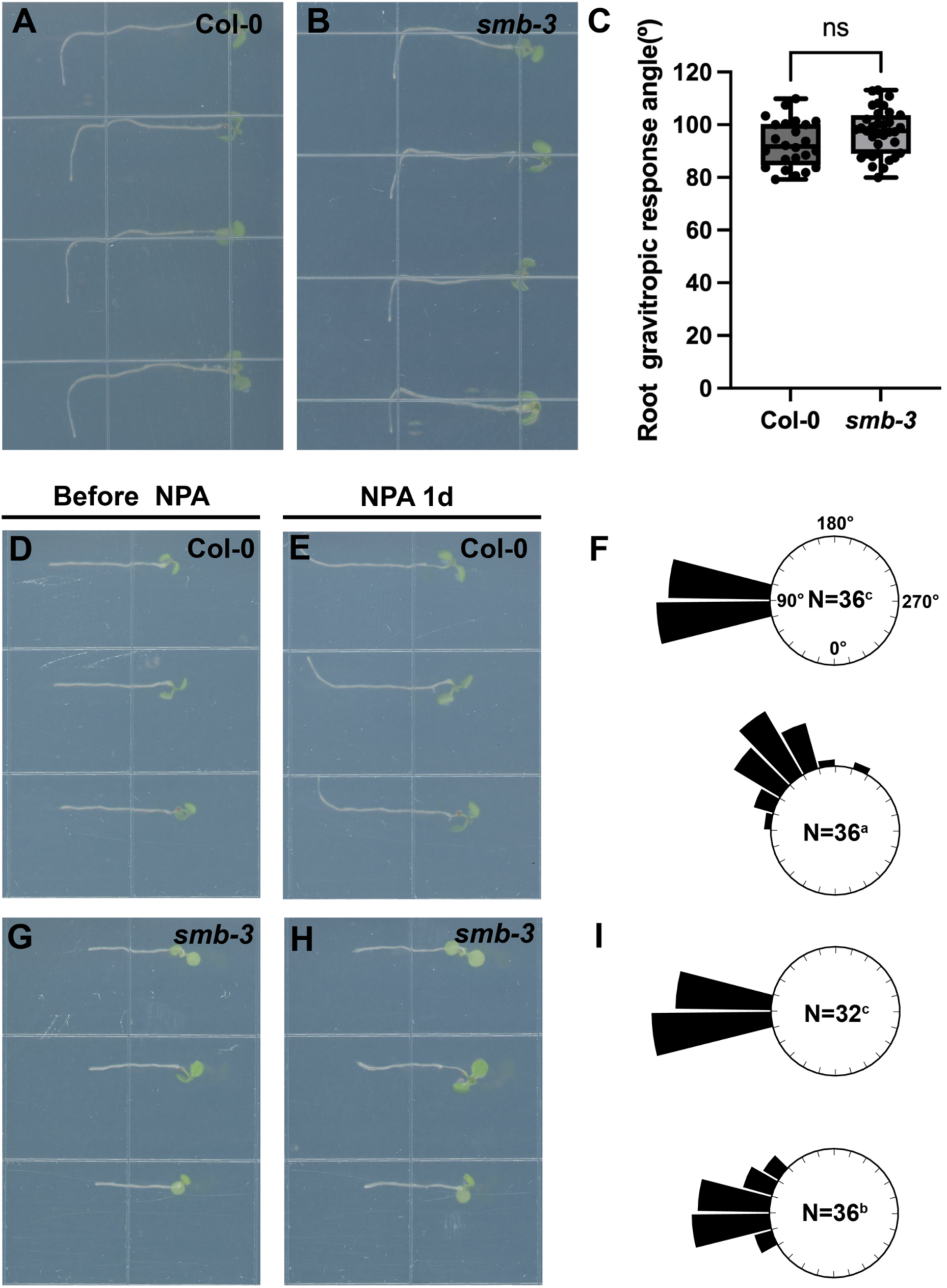
The loss of SMB function does not impact gravitropic root growth. (A and B) Gravitropic root response of Col-0 and *smb-3* mutant. 4-day-old seedlings that grown vertically on 1/2 MS medium transfer to a new 1/2 MS medium and horizontally incubated for 24 hours. (C) Statistical analysis of root gravitropic response curvatures after 24 hours gra-stimulation. The number of seedlings analyzed was 24(Col-0) and 35(*smb-3*), respectively. Statistical significance was calculated by unpaired t test with Welch’s correction. (D-I) Gravitropic root response of Col-0 and *smb-3* under 10μM NPA treatment. 4-day-old seedlings that grown vertically on 1/2 MS medium transfer to a new 1/2 MS medium with 10μM NPA and horizontally incubated for 24 hours. Representative seedling images of Col-0(D and E) and *smb-3* (G and H) before and after gravitropic response. Quantification of frequencies of root gravitropic response curvatures after 24hours gra-stimulation (F and I). The bending angle of each root tip was measured as the absolute value relative to the direction of gravity (g). The frequency was calculated as the proportion of roots falling within 15° intervals relative to the total number of analyzed roots for each line. The bar represents relative frequency. N indicates the number of independent seedlings used in the experiment, and letters indicate significant differences (p<0.05, one-way ANOVA with Tukey’s test).

**Figure S10.**
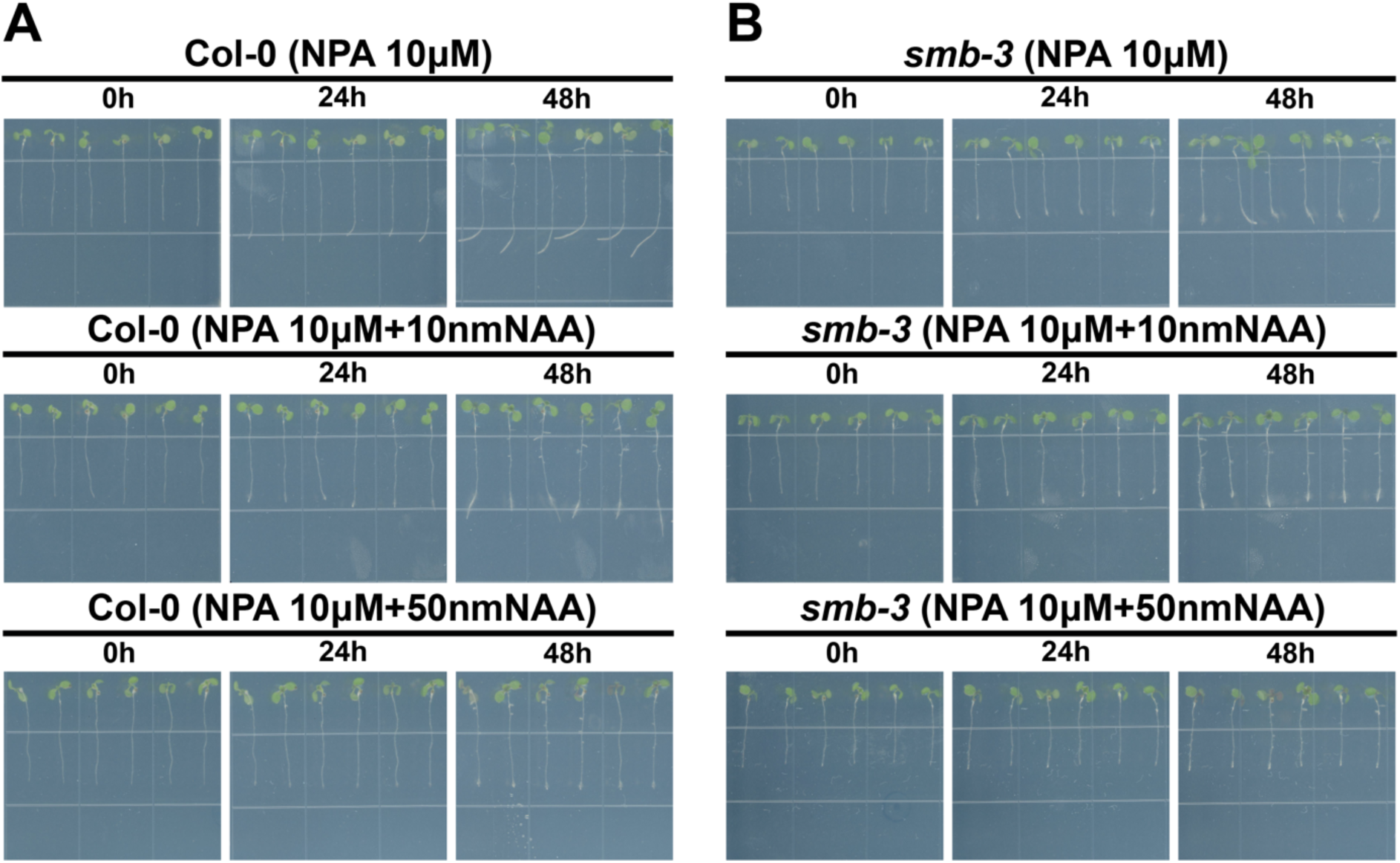
NAA treatment rescues the agravitropic phenotype. (A and B) Time-lapse phenotypic analysis of Col-0(A) and *smb-3*(B) that grown vertically on 1/2 MS medium containing 10μM NPA, with varying concentrations of NAA applied to the root tip.

**Figure S11.**
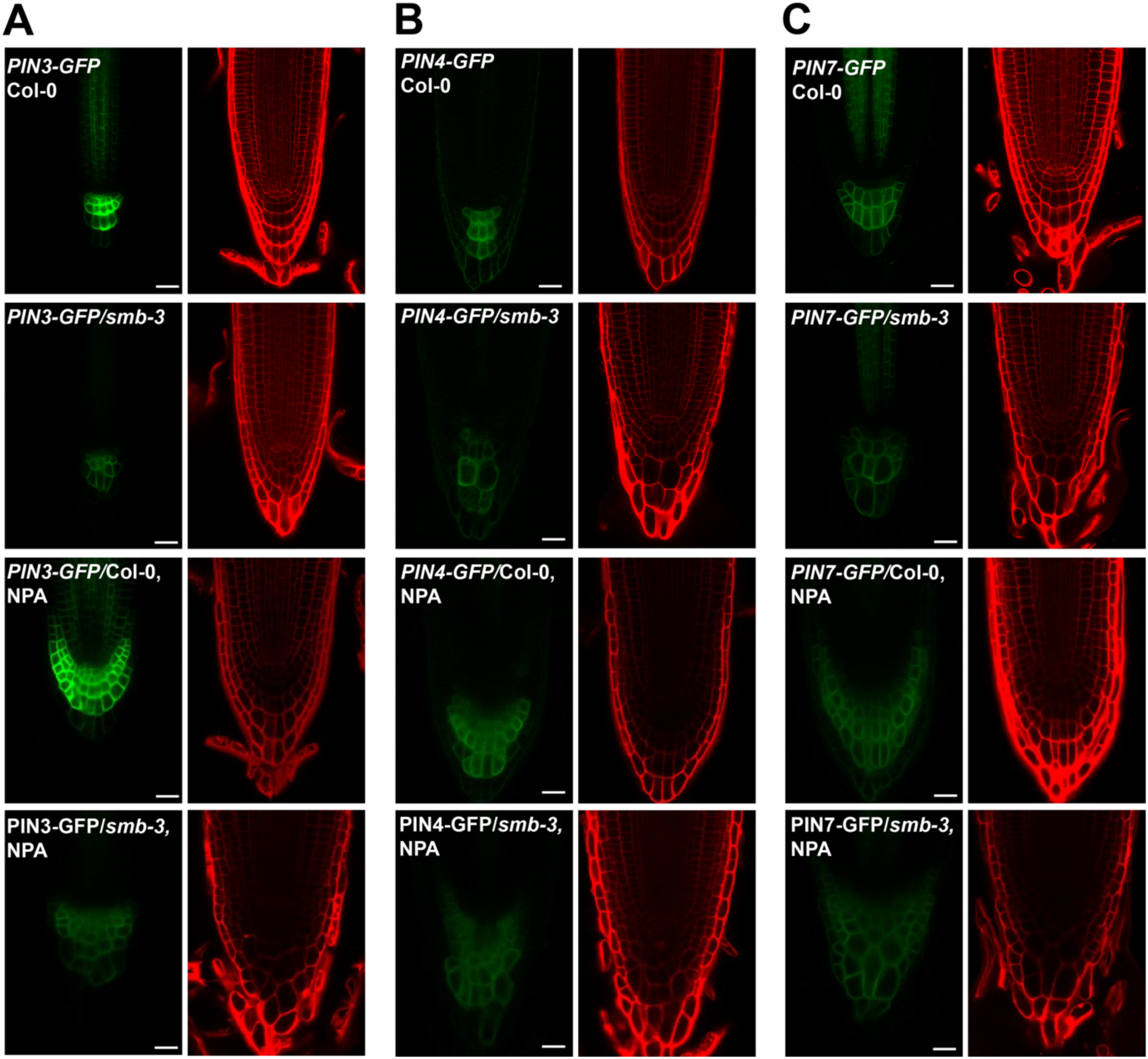
Expression patterns of root cap auxin efflux transporters under NPA treatment. (A-C) Expression patterns of *pPIN3:PIN3-GFP*, *pPIN4:PIN4-GFP* and *pPIN7:PIN7-GFP* in Col-0 and *smb-3* with or without NPA treatment, grown vertically on 1/2 MS medium. Scale bars: 25 μm.

**Figure S12.**
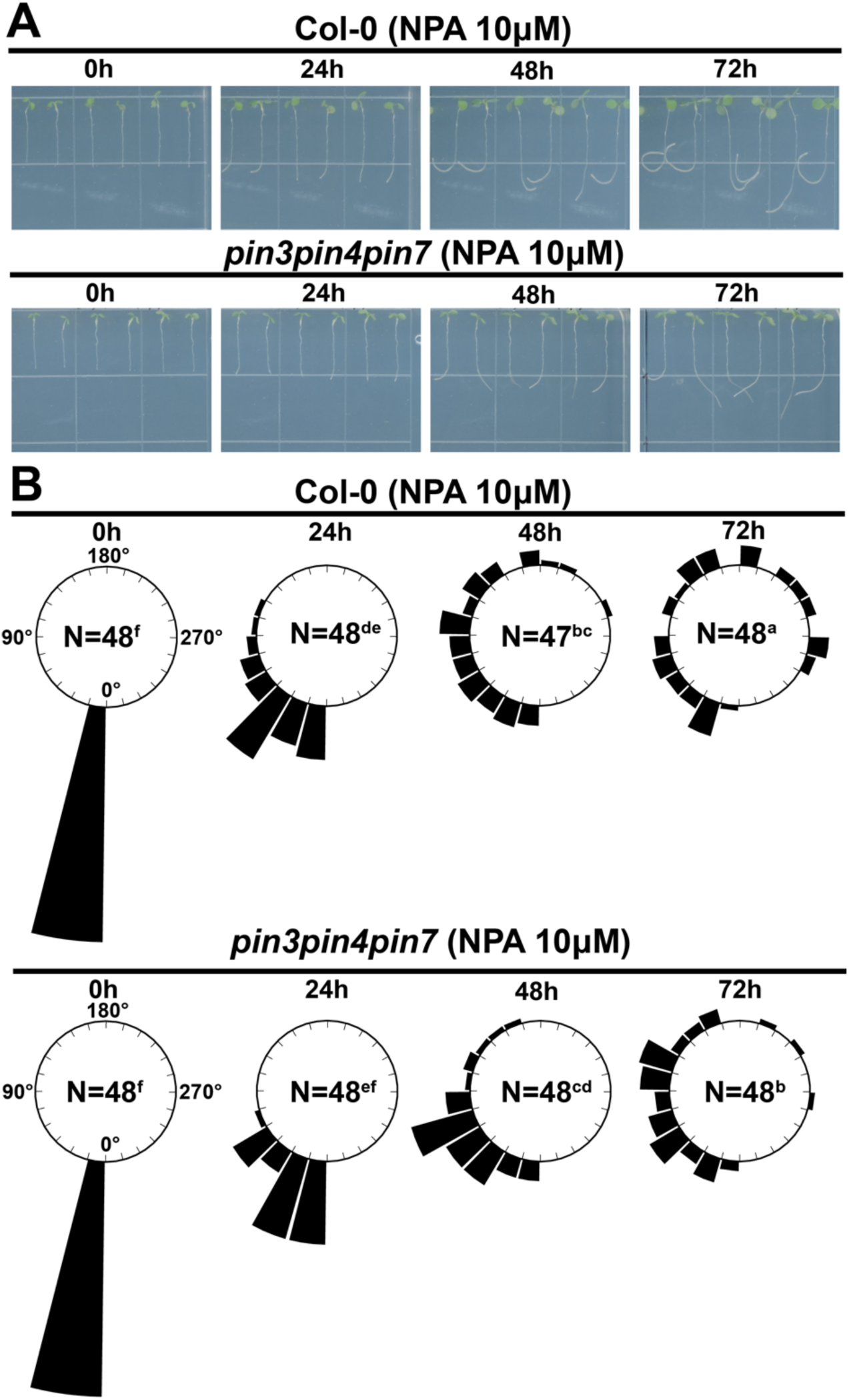
NPA-induced agravitropic growth is compromised in *pin3pin4pin7*. (A) Time-lapse phenotypic analysis of Col-0(upper) and *pin3pin4pin7*(lower) that grown vertically on 1/2 MS medium containing 10μM NPA. (B) Frequencies of root tip growth angles of Col-0(upper) and *pin3pin4pin7*(lower) under NPA treatment. The frequency was calculated as the proportion of roots falling within 15° intervals relative to the total number of analyzed roots for each line. The bar represents relative frequency. N indicates the number of independent seedlings used in the experiment, and letters indicate significant differences (p<0.05, one-way ANOVA with Tukey’s test).

**Figure S13.**
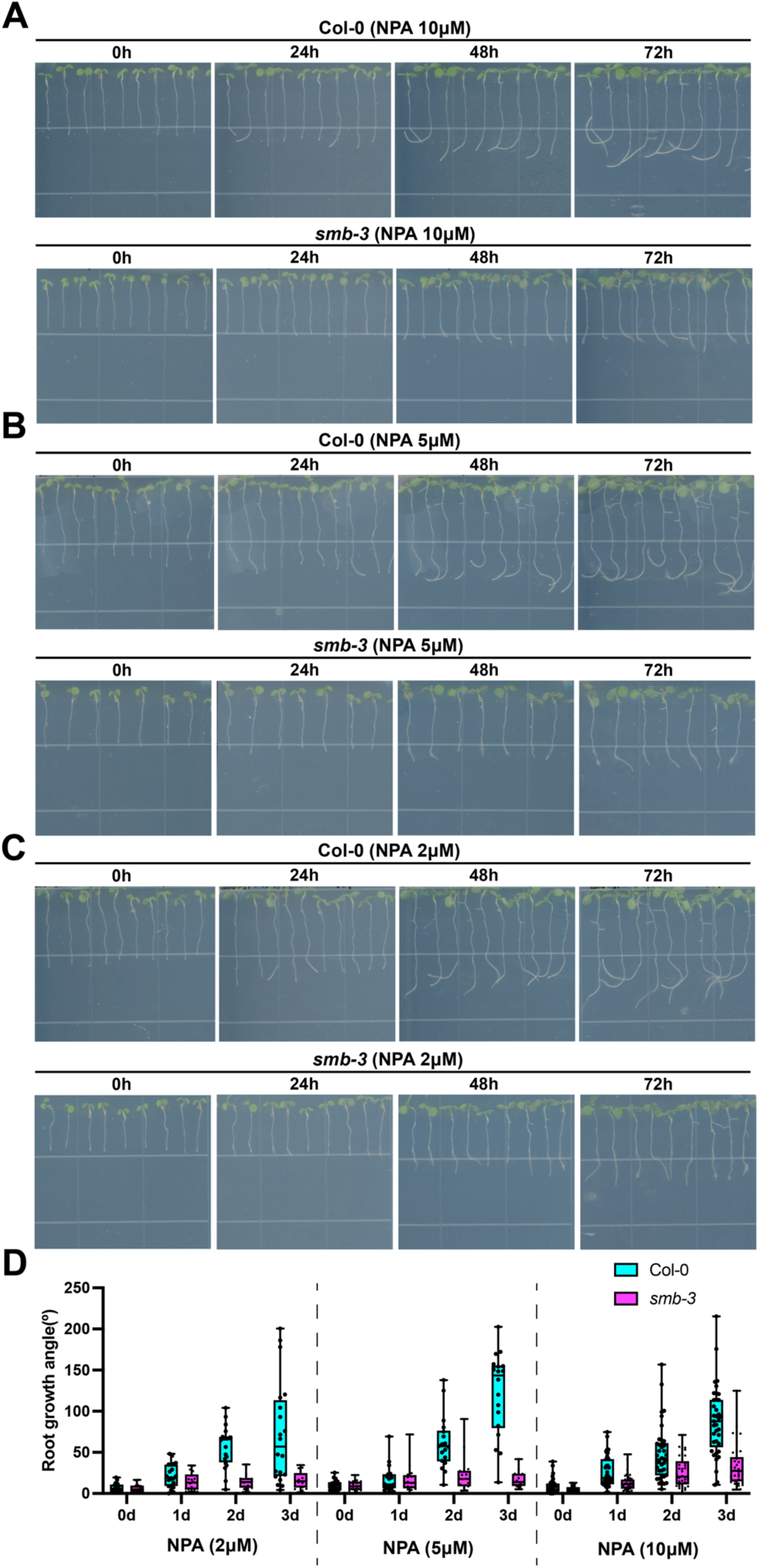
Col-0 displays varying sensitivity to different concentrations of NPA treatment. (A-C) Time-lapse imaging of Col-0 and *smb-3* that grown vertically on 1/2 MS medium with 10μM(A), 5μM(B) and 2μM(C) NPA. (D) Statistical analysis of root tip growth angles with varying concentration of NPA treatment. The direction of each root tip was measured as the absolute value toward the direction of gravity (g).

## REFERENCES

1. Benjamins, R., and Scheres, B. (2008). Auxin: The looping star in plant development. Annual Review of Plant Biology 59, 443–465. 10.1146/annurev.arplant.58.032806.103805.

2. Brumos, J., Robles, L.M., Yun, J., Vu, T.C., Jackson, S., Alonso, J.M., and Stepanova, A.N. (2018). Local Auxin Biosynthesis Is a Key Regulator of Plant Development. Dev Cell 47, 306–318. 10.1016/j.devcel.2018.09.022.

3. Cohen, J.D., and Strader, L.C. (2024). An auxin research odyssey: 1989-2023. Plant Cell 36, 1410–1428. 10.1093/plcell/koae054.

4. Swarup, R., and Bennett, M. (2003). Auxin transport: the fountain of life in plants? Dev Cell 5, 824–826. 10.1016/s1534-5807(03)00370-8.

5. Friml, J., Vieten, A., Sauer, M., Weijers, D., Schwarz, H., Hamann, T., Offringa, R., and Jurgens, G. (2003). Efflux-dependent auxin gradients establish the apical-basal axis of Arabidopsis. Nature 426, 147–153. 10.1038/nature02085.

6. Cheng, Y.F., Dai, X.H., and Zhao, Y.D. (2007). Auxin synthesized by the YUCCA flavin Monooxygenases is essential for embryogenesis and leaf formation in. Plant Cell 19, 2430–2439. 10.1105/tpc.107.053009.

7. Robert, H.S., Grones, P., Stepanova, A.N., Robles, L.M., Lokerse, A.S., Alonso, J.M., Weijers, D., and Friml, J. (2013). Local auxin sources orient the apical-basal axis in Arabidopsis embryos. Curr Biol 23, 2506–2512. 10.1016/j.cub.2013.09.039.

8. Grieneisen, V.A., Xu, J., Marée, A.F.M., Hogeweg, P., and Scheres, B. (2007). Auxin transport is sufficient to generate a maximum and gradient guiding root growth. Nature 449, 1008–1013. 10.1038/nature06215.

9. Rosquete, M.R., Barbez, E., and Kleine-Vehn, J. (2012). Cellular Auxin Homeostasis: Gatekeeping Is Housekeeping. Molecular Plant 5, 772–786. 10.1093/mp/ssr109.

10. Dubreuil, C., Jin, X., Grönlund, A., and Fischer, U. (2018). A local auxin gradient regulates root cap self-renewal and size homeostasis. Current Biology 28, 2581–2587. e2583. 10.1016/j.cub.2018.05.090.

11. Weijers, D., Schlereth, A., Ehrismann, J.S., Schwank, G., Kientz, M., and Jurgens, G. (2006). Auxin triggers transient local signaling for cell specification in Arabidopsis embryogenesis. Dev Cell 10, 265–270. 10.1016/j.devcel.2005.12.001.

12. Noh, B., Murphy, A.S., and Spalding, E.P. (2001). Multidrug resistance-like genes of Arabidopsis required for auxin transport and auxin-mediated development. Plant Cell 13, 2441–2454. 10.1105/tpc.010350.

13. Park, J., Kim, Y.S., Kim, S.G., Jung, J.H., Woo, J.C., and Park, C.M. (2011). Integration of auxin and salt signals by the NAC transcription factor NTM2 during seed germination in Arabidopsis. Plant Physiol 156, 537–549. 10.1104/pp.111.177071.

14. Van Norman, J.M., Xuan, W., Beeckman, T., and Benfey, P.N. (2013). To branch or not to branch: the role of pre-patterning in lateral root formation. Development 140, 4301–4310. 10.1242/dev.090548.

15. Kacprzyk, J., Burke, R., Schwarze, J., and McCabe, P.F. (2022). Plant programmed cell death meets auxin signalling. Febs J 289, 1731–1745. 10.1111/febs.16210.

16. Ganesh, A., Shukla, V., Mohapatra, A., George, A.P., Bhukya, D.P.N., Das, K.K., Kola, V.S.R., Suresh, A., and Ramireddy, E. (2022). Root Cap to Soil Interface: A Driving Force Toward Plant Adaptation and Development. Plant Cell Physiol 63, 1038–1051. 10.1093/pcp/pcac078.

17. Xuan, W., Band, L.R., Kumpf, R.P., Van Damme, D., Parizot, B., De Rop, G., Opdenacker, D., Moller, B.K., Skorzinski, N., Njo, M.F., et al. (2016). Cyclic programmed cell death stimulates hormone signaling and root development in Arabidopsis. Science 351, 384–387. 10.1126/science.aad2776.

18. Michniewicz, M., Brewer, P.B., and Friml, J.I. (2007). Polar auxin transport and asymmetric auxin distribution. Arabidopsis Book 5, e0108. 10.1199/tab.0108.

19. Kleine-Vehn, J., and Friml, J. (2008). Polar targeting and endocytic recycling in auxin-dependent plant development. Annu Rev Cell Dev Biol 24, 447–473. 10.1146/annurev.cellbio.24.110707.175254.

20. Janacek, D.P., Kolb, M., Schulz, L., Mergner, J., Kuster, B., Glanc, M., Friml, J., Ten Tusscher, K., Schwechheimer, C., and Hammes, U.Z. (2024). Transport properties of canonical PIN-FORMED proteins from Arabidopsis and the role of the loop domain in auxin transport. Dev Cell. 10.1016/j.devcel.2024.09.020.

21. Blilou, I., Xu, J., Wildwater, M., Willemsen, V., Paponov, I., Friml, J., Heidstra, R., Aida, M., Palme, K., and Scheres, B. (2005). The PIN auxin efflux facilitator network controls growth and patterning in Arabidopsis roots. Nature 433, 39–44. 10.1038/nature03184.

22. Kleine-Vehn, J., Ding, Z.J., Jones, A.R., Tasaka, M., Morita, M.T., and Friml, J. (2010). Gravity-induced PIN transcytosis for polarization of auxin fluxes in gravity-sensing root cells. P Natl Acad Sci USA 107, 22344–22349. 10.1073/pnas.1013145107.

23. Teale, W., and Palme, K. (2018). Naphthylphthalamic acid and the mechanism of polar auxin transport. J Exp Bot 69, 303–312. 10.1093/jxb/erx323.

24. Abas, L., Kolb, M., Stadlmann, J., Janacek, D.P., Lukic, K., Schwechheimer, C., Sazanov, L.A., Mach, L., Friml, J., and Hammes, U.Z. (2021). Naphthylphthalamic acid associates with and inhibits PIN auxin transporters. Proc Natl Acad Sci U S A 118, e2020857118. 10.1073/pnas.2020857118.

25. Ogura, T., Goeschl, C., Filiault, D., Mirea, M., Slovak, R., Wolhrab, B., Satbhai, S.B., and Busch, W. (2019). Root System Depth in Arabidopsis Is Shaped by via the Dynamic Modulation of Auxin Transport. Cell 178, 400–412. 10.1016/j.cell.2019.06.021.

26. Li, Y.T., Dai, X.H., Cheng, Y.F., and Zhao, Y.D. (2011). Genes Play an Essential Role in Root Gravitropic Responses in Arabidopsis. Molecular Plant 4, 171–179. 10.1093/mp/ssq052.

27. Stepanova, A.N., Robertson-Hoyt, J., Yun, J., Benavente, L.M., Xie, D.Y., DoleZal, K., Schlereth, A., Jürgens, G., and Alonso, J.M. (2008). TAA1-mediated auxin biosynthesis is essential for hormone crosstalk and plant development. Cell 133, 177–191. 10.1016/j.cell.2008.01.047.

28. Ma, W.Y., Li, J.J., Qu, B.Y., He, X., Zhao, X.Q., Li, B., Fu, X.D., and Tong, Y.P. (2014). Auxin biosynthetic gene is involved in low nitrogen-mediated reprogramming of root architecture in Arabidopsis. Plant J 78, 70–79. 10.1111/tpj.12448.

29. Wang, J.-W., Wang, L.-J., Mao, Y.-B., Cai, W.-J., Xue, H.-W., and Chen, X.-Y. (2005). Control of root cap formation by microRNA-targeted auxin response factors in Arabidopsis. The Plant Cell 17, 2204–2216. 10.1105/tpc.105.033076.

30. Lv, B., Yu, Q., Liu, J., Wen, X., Yan, Z., Hu, K., Li, H., Kong, X., Li, C., Tian, H., et al. (2020). Non-canonical AUX/IAA protein IAA33 competes with canonical AUX/IAA repressor IAA5 to negatively regulate auxin signaling. EMBO J 39, e101515. 10.15252/embj.2019101515.

31. Cai, X., Zhang, H., Mu, C., Chen, Y., He, C., Liu, M., Laux, T., and Pi, L. (2024). A mobile miR160-triggered transcriptional axis controls root stem cell niche maintenance and regeneration in Arabidopsis. Dev Cell 60, 1–13. 10.1016/j.devcel.2024.10.006.

32. Zheng, L., Hu, Y., Yang, T., Wang, Z., Wang, D., Jia, L., Xie, Y., Luo, L., Qi, W., Lv, Y., et al. (2024). A root cap-localized NAC transcription factor controls root halotropic response to salt stress in Arabidopsis. Nat Commun 15, 2061. 10.1038/s41467-024-46482-7.

33. Willemsen, V., Bauch, M., Bennett, T., Campilho, A., Wolkenfelt, H., Xu, J., Haseloff, J., and Scheres, B. (2008). The NAC domain transcription factors FEZ and SOMBRERO control the orientation of cell division plane in Arabidopsis root stem cells. Dev Cell 15, 913–922. 10.1016/j.devcel.2008.09.019.

34. Kamiya, M., Higashio, S.-Y., Isomoto, A., Kim, J.-M., Seki, M., Miyashima, S., and Nakajima, K. (2016). Control of root cap maturation and cell detachment by BEARSKIN transcription factors in Arabidopsis. Development 143, 4063–4072. 10.1242/dev.142331.

35. Bennett, T., van den Toorn, A., Sanchez-Perez, G.F., Campilho, A., Willemsen, V., Snel, B., and Scheres, B. (2010). SOMBRERO, BEARSKIN1, and BEARSKIN2 regulate root cap maturation in Arabidopsis. The Plant Cell 22, 640–654. 10.1105/tpc.109.072272.

36. Ravelo-Ortega, G., Pelagio-Flores, R., López-Bucio, J., Campos-García, J., de la Cruz, H.R., and López-Bucio, J.S. (2022). Early sensing of phosphate deprivation triggers the formation of extra root cap cell layers via SOMBRERO through a process antagonized by auxin signaling. Plant Mol Biol 108, 77–91. 10.1007/s11103-021-01224-x.

37. Huysmans, M., Buono, R.A., Skorzinski, N., Radio, M.C., De Winter, F., Parizot, B., Mertens, J., Karimi, M., Fendrych, M., and Nowack, M.K. (2018). NAC Transcription Factors ANAC087 and ANAC046 Control Distinct Aspects of Programmed Cell Death in the Arabidopsis Columella and Lateral Root Cap. Plant Cell 30, 2197–2213. 10.1105/tpc.18.00293.

38. Fendrych, M., Van Hautegem, T., Van Durme, M., Olvera-Carrillo, Y., Huysmans, M., Karimi, M., Lippens, S., Guerin, C.J., Krebs, M., Schumacher, K., and Nowack, M.K. (2014). Programmed cell death controlled by ANAC033/SOMBRERO determines root cap organ size in Arabidopsis. Curr Biol 24, 931–940. 10.1016/j.cub.2014.03.025.

39. Wang, J., Bollier, N., Buono, R.A., Vahldick, H., Lin, Z., Feng, Q., Hudecek, R., Jiang, Q., Mylle, E., Van Damme, D., and Nowack, M.K. (2024). A developmentally controlled cellular decompartmentalization process executes programmed cell death in the Arabidopsis root cap. Plant Cell 36, 941–962. 10.1093/plcell/koad308.

40. Kang, B.H., Anderson, C.T., Arimura, S.I., Bayer, E., Bezanilla, M., Botella, M.A., Brandizzi, F., Burch-Smith, T.M., Chapman, K.D., Dunser, K., et al. (2022). A glossary of plant cell structures: Current insights and future questions. Plant Cell 34, 10–52. 10.1093/plcell/koab247.

41. Staehelin, L.A., Giddings, T.H., Kiss, J.Z., and Sack, F.D. (1990). Macromolecular Differentiation of Golgi Stacks in Root-Tips of Arabidopsis and Nicotiana Seedlings as Visualized in High-Pressure Frozen and Freeze-Substituted Samples. Protoplasma 157, 75–91. 10.1007/Bf01322640.

42. Wang, P., Liang, Z., and Kang, B.H. (2019). Electron tomography of plant organelles and the outlook for correlative microscopic approaches. New Phytol 223, 1756–1761. 10.1111/nph.15882.

43. Liu, Z., Wang, P., Goh, T., Nakajima, K., and Kang, B.H. (2024). Mucilage secretion from the root cap requires the NAC family transcription factor BEARSKIN2. Plant Physiol 196, 1180–1195. 10.1093/plphys/kiae402.

44. Vicre, M., Santaella, C., Blanchet, S., Gateau, A., and Driouich, A. (2005). Root border-like cells of Arabidopsis. Microscopical characterization and role in the interaction with rhizobacteria. Plant Physiol 138, 998–1008. 10.1104/pp.104.051813.

45. Wang, P.F., Chen, X.S., Goldbeck, C., Chung, E., and Kang, B.H. (2017). A distinct class of vesicles derived from the trans-Golgi mediates secretion of xylogalacturonan in the root border cell. Plant J 92, 596–610. 10.1111/tpj.13704.

46. Wang, P., and Kang, B.H. (2018). The trans-Golgi sorting and the exocytosis of xylogalacturonan from the root border/border-like cell are conserved among monocot and dicot plant species. Plant Signal Behav 13, e1469362. 10.1080/15592324.2018.1469362.

47. O’Malley, R.C., Huang, S.C., Song, L., Lewsey, M.G., Bartlett, A., Nery, J.R., Galli, M., Gallavotti, A., and Ecker, J.R. (2016). Cistrome and Epicistrome Features Shape the Regulatory DNA Landscape. Cell 165, 1280–1292. 10.1016/j.cell.2016.04.038.

48. Zemlyanskaya, E.V., Omelyanchuk, N.A., Ubogoeva, E.V., and Mironova, V.V. (2018). Deciphering Auxin-Ethylene Crosstalk at a Systems Level. International Journal of Molecular Sciences 19, 4060. 10.3390/ijms19124060.

49. Di Mambro, R., De Ruvo, M., Pacifici, E., Salvi, E., Sozzani, R., Benfey, P.N., Busch, W., Novak, O., Ljung, K., Di Paola, L., et al. (2017). Auxin minimum triggers the developmental switch from cell division to cell differentiation in the Arabidopsis root. Proc Natl Acad Sci U S A 114, E7641–E7649.10.1073/pnas.1705833114.

50. Pierdonati, E., Unterholzner, S.J., Salvi, E., Svolacchia, N., Bertolotti, G., Dello Ioio, R., Sabatini, S., and Di Mambro, R. (2019). Cytokinin-Dependent Control of GH3 Group II Family Genes in the Arabidopsis Root. Plants 8, 94. 10.3390/plants8040094.

51. Kumpf, R.P., and Nowack, M.K. (2015). The root cap: a short story of life and death. J Exp Bot 66, 5651–5662. 10.1093/jxb/erv295.

52. Nicolas, W.J., Grison, M.S., Trepout, S., Gaston, A., Fouche, M., Cordelieres, F.P., Oparka, K., Tilsner, J., Brocard, L., and Bayer, E.M. (2017). Architecture and permeability of post-cytokinesis plasmodesmata lacking cytoplasmic sleeves. Nat Plants 3, 17082. 10.1038/nplants.2017.82.

53. Burch-Smith, T.M., and Zambryski, P.C. (2012). Plasmodesmata Paradigm Shift: Regulation from Without Versus Within. Annual Review of Plant Biology 63, 239–260. 10.1146/annurev-arplant-042811-105453.

54. Han, X., Hyun, T.K., Zhang, M.H., Kumar, R., Koh, E.J., Kang, B.H., Lucas, W.J., and Kim, J.Y. (2014). Auxin-Callose-Mediated Plasmodesmal Gating Is Essential for Tropic Auxin Gradient Formation and Signaling. Dev Cell 28, 132–146. 10.1016/j.devcel.2013.12.008.

55. Li, M., Wang, M., Lin, Q., Wang, M., Niu, X., Cheng, J., Xu, M., Qin, Y., Liao, X., Xu, J., and Wu, S. (2022). Symplastic communication in the root cap directs auxin distribution to modulate root development. J Integr Plant Biol 64, 859–870. 10.1111/jipb.13237.

56. Di Mambro, R., Svolacchia, N., Dello Ioio, R., Pierdonati, E., Salvi, E., Pedrazzini, E., Vitale, A., Perilli, S., Sozzani, R., Benfey, P.N., et al. (2019). The Lateral Root Cap Acts as an Auxin Sink that Controls Meristem Size. Current Biology 29, 1199–1205. 10.1016/j.cub.2019.02.022.

57. Bennett, T., Hines, G., van Rongen, M., Waldie, T., Sawchuk, M.G., Scarpella, E., Ljung, K., and Leyser, O. (2016). Connective Auxin Transport in the Shoot Facilitates Communication between Shoot Apices. PLoS Biol 14, e1002446. 10.1371/journal.pbio.1002446.

